# Loss of UFMylation supports prostate cancer metastasis and rewires cell metabolism towards hexosamine biosynthesis

**DOI:** 10.1101/2025.06.02.657324

**Authors:** Laura Bozal-Basterra, María-Camila Salazar, Ana Margarida Ferreira Campos, Margherita Demicco, Daniel R. Schmidt, Klaudia Sobczak, June Ereño-Orbea, Patricia Altea-Manzano, Ainara Miranda Villanueva, Belén Martínez La Osa, Saioa Garcia-Longarte, María Ponce-Rodriguez, Isabel Mendizabal, Onintza Carlevaris, Ianire Astobiza, Natalia Martin-Martin, Amaia Zabala-Letona, Ana Talamillo, Juan Fernández-García, Mikel Azkargorta, Ibon Iloro, Rosa Barrio, Félix Elortza, Sarah-Maria Fendt, Matthew G. Vander Heiden, Jesús-Jiménez Barbero, James D. Sutherland, Arkaitz Carracedo

## Abstract

The acquisition of metastatic features in tumor cells encompasses genetic and non-genetic adaptation, including reprogramming of cellular metabolism. Here we show that loss of UFMylation reroutes glucose metabolism, promotes invasive capacity and supports prostate cancer metastasis. Through transcriptome-based bioinformatics analysis, we identified a reduction in the ubiquitin-like modifier UFM1 and its ligase UFL1 in metastatic prostate cancer. We demonstrate that loss of UFMylation results in enhanced cancer cell dissemination and a switch from cellular proliferation to invasion. Using biotin-based proteomics, we identified phosphofructokinase (PFKAP) as an unprecedented UFMylation substrate. Consistent with UFMylation playing a role in the regulation of phosphofructokinase activity, loss of UFMylation reduced glucose metabolism in favour of hexosamine biosynthesis, which resulted in elevated glycosylation of proteins relevant for cell invasion. These results reveal a role for UFMylation in the regulation of phosphofructokinase and glucose metabolism to support prostate cancer metastasis.

---

Throughout the process of cancer progression, specific molecular and metabolic programs support the survival, growth and dissemination of tumor cells^1^.To identify genes and processes implicated in prostate cancer metastasis, we analysed transcriptomic datasets that contained data from normal, primary and metastatic prostate cancer specimens, as well as clinical follow-up. We focused our attention on genes whose expression was selectively altered in metastatic specimens, but not in primary tumors regardless of the prognosis (**Fig. 1A and Supplementary Table 1**). Functional enrichment analysis to identify genes with selectively altered expression in metastasis highlighted a significant alteration of protein UFMylation^2^ in 2 different gene ontology categories (**Fig. 1B and Supplementary Table 2).** UFMylation-related genes encoding for the covalently bound ubiquitin-like protein UFM1 and its ligase UFL1 were downregulated in prostate metastatic specimens from various datasets (**Fig. 1C, Extended Data Fig. 1A and Supplementary Table 1**), a finding consistent with a high frequency of deletions in multiple prostate cancer patient cohorts (**Fig. 1D, Extended Data Fig. 1B-C**).

**Fig. 1.**
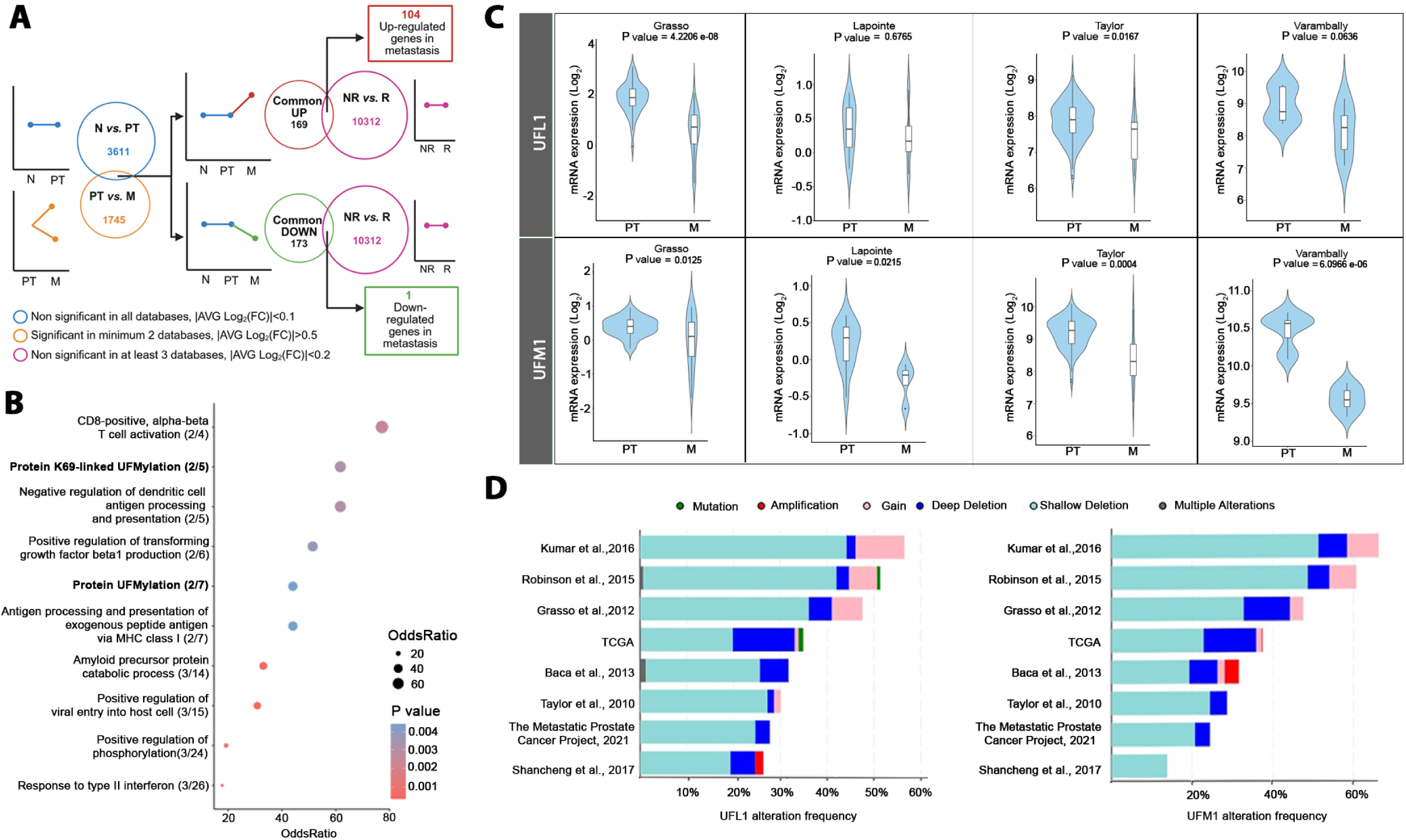
*UFL1* and *UFM1*, core genes in the process of UFMylation, are downregulated in metastatic prostate cancer. **A.** Schematic overview of the selection criteria to identify genes altered in metastasis based on at least 4 transcriptomic databases of human prostate cancer specimens. The Log_2_(fold change) is calculated with limma package. The p value is obtained from a limma differential expression analysis between the two groups. In brief, genes selected complied with 3 criteria: i) non-significant differences (p value > 0.05) between normal (N) and primary tumor (PT) in any of the datasets (blue circle) (Grasso, Lapointe, Taylor, Tomlins and Varambally), ii) significant differences (p value < 0.05) between PT and metastasis (M) in at least 2 different datasets (orange circle), and iii) non-significant differences (p value > 0.05) between non-recurrent (NR) and recurrent (R) patients in at least 3 different datasets (pink circle). Genes were classified as upregulated (95) or downregulated (119) in metastasis. Threshold of the average fold change (FC) in gene expression in the datasets where the gene appears: N *vs.* PT: |AVG Log_2_(FC)| < 0.1; PT *vs*. M: |AVG Log_2_(FC)| > 0.5; NOR *vs.* R: |AVG Log_2_(FC)| < 0.2. **B.** Enrichment results for the 119 downregulated genes in metastasis. Ratio in between parenthesis following ontology terms, shows the number of genes found in our analysis *vs.* the number of genes that the specific gene ontology term contains. Dot size represents the percentage of genes found in our analysis with respect to the number of genes that the specific gene ontology term contains (OddsRatio). Colors represent p value according to a gradient. **C.** Violin plots depicting the expression of *UFL1* (upper panels) and *UFM1* (lower panels) in primary tumor (PT) and metastatic (M) human prostate cancer specimens in four representative datasets (Grasso, Lapointe, Taylor and Varambally). The y-axis represents the Log_2_-normalized gene expression (fluorescence intensity values for microarray data or, sequencing reads values obtained after gene quantification with RSEM and normalization using UpperQuartile in case of RNA-seq). P value derives from unpaired t-test analysis between the indicated groups. *Source:* cancertool.org. **D.** Percentage of prostate cancer patients (X axis) with copy number alterations (CNA) in *UFL1* (left) and *UFM1* (right) genes: mutations in green; amplifications in red; gain in light pink; deep deletions in blue; shallow deletions in aquamarine; multiple alterations in grey in different datasets (Y axis). Deep deletion indicates a deep loss, possibly a homozygous deletion; shallow deletion indicates a shallow loss, possibly a heterozygous deletion; gain indicates a low-level gain (a few additional copies, often broad); amplification indicate a high-level amplification (more copies, often focal). *Source:* cbioportal.org.

UFMylation represents the most recently discovered ubiquitin-like post-translational modification^2^. Similar to ubiquitin, UFM1 covalently binds to its target proteins in a biochemical process involving three different (E1, E2 and E3-like) enzymes (**Extended Data** Fig. 1D)^3–5^. UFMylation plays crucial roles in endoplasmic reticulum homeostasis^6^, genomic instability^7–9^, protein synthesis^10,11^, erythroid^12^ and liver development^13^, immune system^14^, cancer^15,16^ and other cellular processes^17,18^. To date, only a few substrates of UFMylation have been reported^19,20^, and the biological significance of UFMylation in cancer progression remains poorly understood. To test the role of UFMylation in prostate cancer biology, we silenced *UFL1* and *UFM1* in PC3 metastatic prostate cancer cells *in vitro* (**Extended Data Fig. 2A**). Interestingly, both *UFL1*- (shUFL1 #1 and shUFL1 #2) and *UFM1*-silenced cells (shUFM1 #1) showed impaired cell growth and colony formation capacity (**Fig. 2A**, **Extended Data Fig. 2B-C**). In contrast, *UFL1* and *UFM1* silencing increased the invasive capacity of PC3 cells in a collagen I: Matrigel matrix, and in spheroid assays using a collagen matrix, as well as in Matrigel-coated transwell assays (**Fig. 2B-C, Extended Data Fig. 2D-E**). Thus, these *in vitro* assays suggest an increase of invasive properties at the expense of cell proliferation upon inhibition of the UFMylation machinery. These changes in proliferation and invasion upon *UFL1* silencing (**Extended Data Fig. 2F)** were recapitulated in two additional metastatic prostate cancer cell lines, DU145 and 22RV1 (**Extended Data Fig. 2G-H)**. To ascertain the net outcome of biological reprogramming elicited by silencing of the UFMylation machinery, we inoculated bioluminescent control (shScramble or shSc) or *UFL1*-silenced prostate cancer cells (shUFL1 #1) intracardially and monitored disseminated tumor growth in nude mice. In line with the reduction in gene expression and gene copy number observed in metastatic prostate cancer specimens, mice injected with *UFL1*-silenced PC3 cells exhibited increased dissemination observed by bioluminescence imaging (**Fig. 2D-E**), and displayed evidence of significantly higher metastasis in bones (**Fig. 2F-G**) and soft tissues, such as the lungs (**Extended Data Fig. 2I)**. Taken together, these data reveal that experimental reduction of the UFMylation machinery elicits a switch from cell proliferation to invasion that sustains metastatic dissemination.

**Fig. 2.**
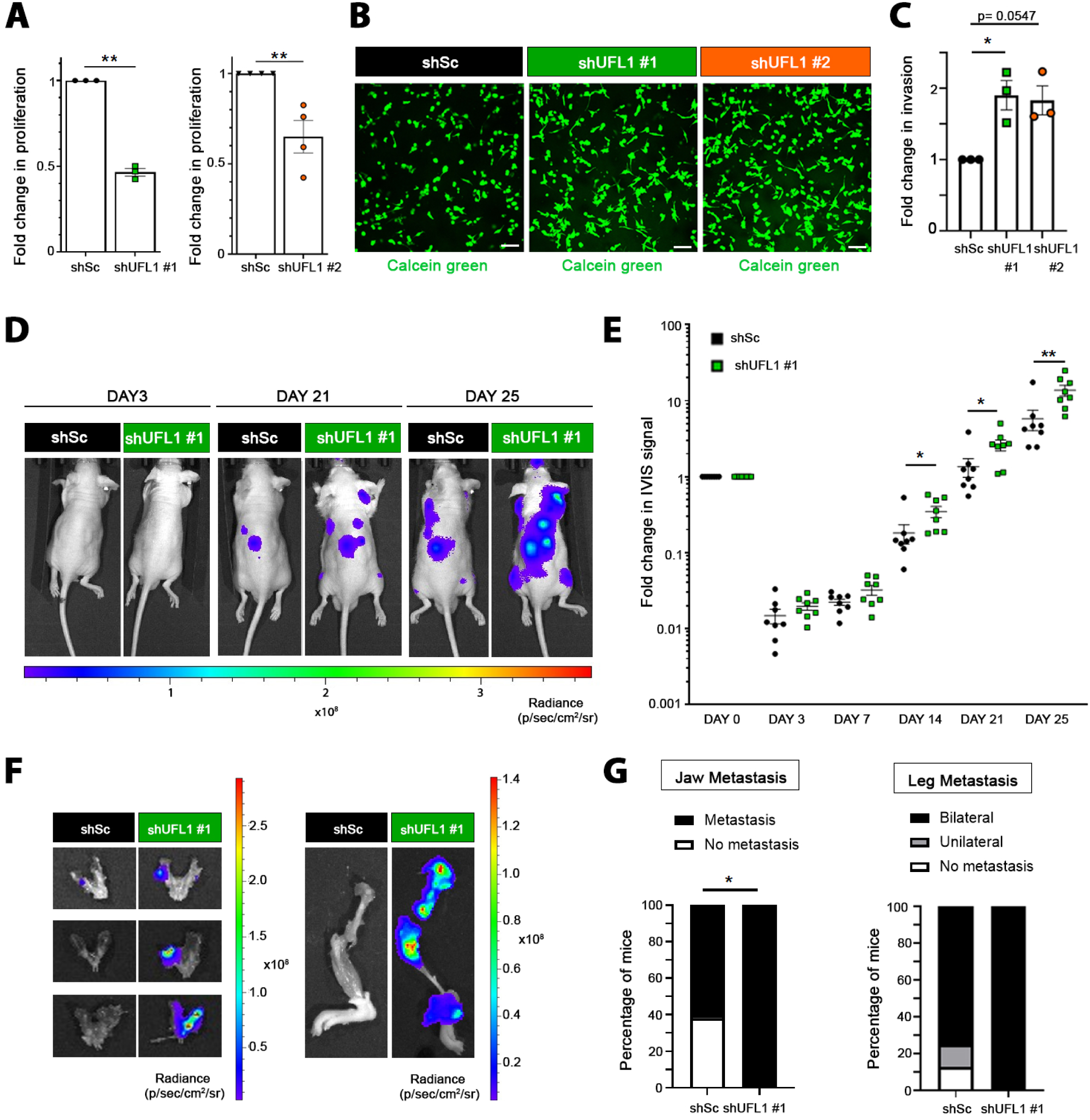
Reduced *UFL1* levels promote a switch from proliferation to invasion and support metastatic dissemination. **A.** Analysis of cell proliferation by crystal violet staining of control (shSc) and *UFL1*-silenced (shUFL1 #1 and shUFL1 #2) PC3 cells. The fold change in crystal violet absorbance at 6 days normalized to shSc is represented. P value was obtained by one-sample t-test. shSc *vs.* shUFL1 #1 (n=3); shSc *vs.* shUFL1 #2 (n=4). **B.** Representative images of the invasive ability of control (shSc) and *UFL1*-silenced (shUFL1 #1 and shUFL1 #2) PC3 cells in a collagen:matrigel 3D matrix. The invasive area of cancer cells was visualized by calcein green staining. Scale bar: 100 µm. **C.** Quantification of the invasive ability of control (shSc) and *UFL1*-silenced (shUFL1 #1 and shUFL1 #2) PC3 cells in a collagen:matrigel 3D matrix. The total area occupied by calcein green-stained cells was quantified using ImageJ. Each dot represents a different experiment (n=3). P value was obtained by ordinary one-way ANOVA with multiple comparison analysis. **D-E.** IVIS relative flux data along the experimental process: PC3 GFP LUC cells transduced with control (shSc) or shUFL1 #1 were injected intracardiacally in nude mice and followed up to 25 days. Representative images (D) and the whole body photon flux normalized to time 0 (E) are represented. Each dot represents a different mouse (n=8 mice per group). P value was obtained by Mann Whitney tests. **F-G.** *Ex vivo* jaw and leg metastasis from mice in (D-E). (F) Representative images of jaws (left) and legs (right) from 3 different mice per group (2 groups: mice injected with shSc and mice injected with shUFL1 #1 PC3 cells) are represented. (G) Contingency analysis of metastatic lesions (n=8 mice per group) in jaws (left) and in legs (right). For jaw metastasis, one-tailed Chi-square test was applied for statistical analysis.

UFMylation regulates a range of biological processes^6–20^. We reasoned that the increased invasive and metastatic capacity of cells with compromised UFMylation results from reduced conjugation of UFM1 onto specific target proteins. To identify candidate UFMylated proteins, we carried out proximity proteomics using UFM1 as bait (TurboID^21^, TbID-UFM1) and UFM1 conjugation analysis using bioUFM1^22^ (**Fig. 3A**). Total lysates from prostate cancer cells expressing TbID-UFM1 (*vs*. cells expressing the TbID moiety alone) and bioUFM1 (*vs*. cells expressing the BirA moiety alone) were subjected to streptavidin pulldown and analysed by liquid chromatography tandem mass spectrometry (LC-MS/MS). We identified known UFM1-conjugating proteins such as the E1 enzyme UBA5, as well as other UFMylation components such as DDRGK domain-containing protein 1 (DDRGK1; also known as UFM1-binding protein 1 (UFBP1)) ^6^ in these experiments, validating our approach (**Extended Data Fig. 3A-B, Supplementary Tables 3-4**). Together with UBA5, only one additional protein was consistently detected and validated in both experimental approaches, namely PFKAP (ATP-dependent 6-phosphofructokinase, platelet type) (**Fig. 3B, Extended Data Fig. 3A-C, Supplementary Tables 3-4)**. Pulldown of tagged PFKAP confirmed this enzyme as a target of UFMylation (**Fig. 3C, Extended Data Fig. 3D**). Of note, a mutant form of UFM1 lacking the C-terminal glycine 83 residue (that cannot be conjugated with target proteins^2^; HA-UFM1 ΔG) could not be detected bound to PFKAP-GFP in the GFP pulldowns, further supporting that UFM1 is enzymatically conjugated to PFKAP (**Fig. 3C)**. Consistent with these data, *UFL1* knockdown using 2 different shRNAs (shUFL1 #1 and shUFL1 #2) decreased the signal of UFM1 in PFKAP pulldowns (**Extended Data Fig. 3E**). To identify the UFMylated residues in PFKAP, we performed mass spectrometry analysis of GFP pulldowns in PFKAP-GFP transfected HEK293FT cells in the presence of HA-UFM1. Mass spectrometry analysis detected five different lysine residues (K) in PFKAP as potentially UFMylated: K281, K287, K395, K625 and K736 (**Extended Data Fig. 3F and Supplementary Table 5).** PFKAP is ubiquitinated on lysine K281^23^ and UFMylation can occur on lysines that are ubiquitinated^14,15^. However, mutating lysine 281-to-arginine (K281R) did not reduce PFKAP UFMylation (**Extended Data Fig. 3G)**, suggesting that other identified lysine residues must be UFMylated in PFKAP. To test this notion, we generated lysine-to-arginine (KR) mutations in all five PFKAP lysine residues identified as UFMylated by mass spectrometry (PFKAP 5KR) and found that this mutant enzyme exhibited significantly reduced UFMylation signal (**Fig. 3D-E**). Based on these analyses, we concluded that PFKAP can be modified by UFM1 on multiple lysines, although it remains possible that additional residues on PFKAP are subject to this covalent modification. PFKAP catalyses the phosphorylation of fructose 6-phosphate (F6P) to fructose 1,6-bisphosphate, an important commitment step in glycolysis. Carbohydrate carbons upstream of PFKAP can have alternative metabolic fates, such as entry into the hexosamine biosynthesis pathway that produces intermediates for protein glycosylation (**Fig. 4A**). Therefore, we hypothesized that altered PFKAP UFMylation influences the fate of carbohydrates regulating the alternative flux from glycolysis to the hexosamine biosynthesis pathway, which would be compatible with a shift in metabolism away from anabolism towards a gain of invasive properties^24,25^. To test this hypothesis, we measured extracellular acidification rate (ECAR), as a surrogate for glycolysis, and oxygen consumption rate (OCR) in *UFL1*-silenced cells (**Fig. 4B**). *UFL1* silencing resulted in lower glycolysis, glycolytic capacity, glycolytic reserve and basal OCR when compared to control cells (shSc) (**Fig. 4B and Extended data Fig. 4A-D**). These data are consistent with the drop in proliferation observed in *UFL1*-silenced prostate cancer cells (**Fig. 2A and Extended data Fig. 2G**). Next, we determined the activity of the hexosamine biosynthesis pathway in *UFL1*-silenced cells using kinetic ^13^C_6_-glucose tracing analysis^26^. We found increased carbon flux through the hexosamine biosynthesis pathway, demonstrated by increased glucose contribution to sialic acid (see M+9 and M+11), CMP-sialic acid (see M+11, M+14 and M+16) and UDP-GlcNAC (see M+13), in *UFL1*-silenced cells (**Fig. 4C, Extended Data Fig. 4E and Supplementary Table 6**). Protein glycosylation is a post-translational modification that governs many cellular processes, and aberrant glycosylation contributes to tumour progression in prostate cancer^27–30^. To test if elevated production of glycosylation intermediates upon reduced UFMylation influences protein glycosylation (sialylation), we monitored β-1,4-N-acetylglucosamine- and sialic-acid-linked proteins using fluorescent wheat germ agglutinin (WGA) confocal imaging in basal conditions (**Fig. 4D**) or after transient inhibition of N-glycosylation in PC3 cells using tunicamycin (0.05 μg ml−1, 72 h and washout for 7 h)^31^ (**Fig. 4E**), which affects the first step of biosynthesis of N-linked glycans^32,33^. We found that *UFL1*-silenced cells exhibited elevated abundance of β-1,4-N-acetylglucosamine- and sialic-acid-linked proteins (**Fig. 4D**), and higher recovery rate of protein glycosylation after transient inhibition of N-glycosylation using tunicamycin (**Fig. 4E**). Furthermore, assessment of α(2,6)- and α(2,3)-linked sialic acids through their binding to lectins SNA (α2,6) or SiaFind α2,3-specific Lectenz (α2,3) confirmed that *UFL1*-silenced cells contained more α(2,6) and α(2,3) sialoglycans by FACS analysis (**Fig. 4F-G**) and more α(2,6) sialic acid-conjugated proteins than control cells by western blot (**Fig. 4H**).

**Fig. 3.**
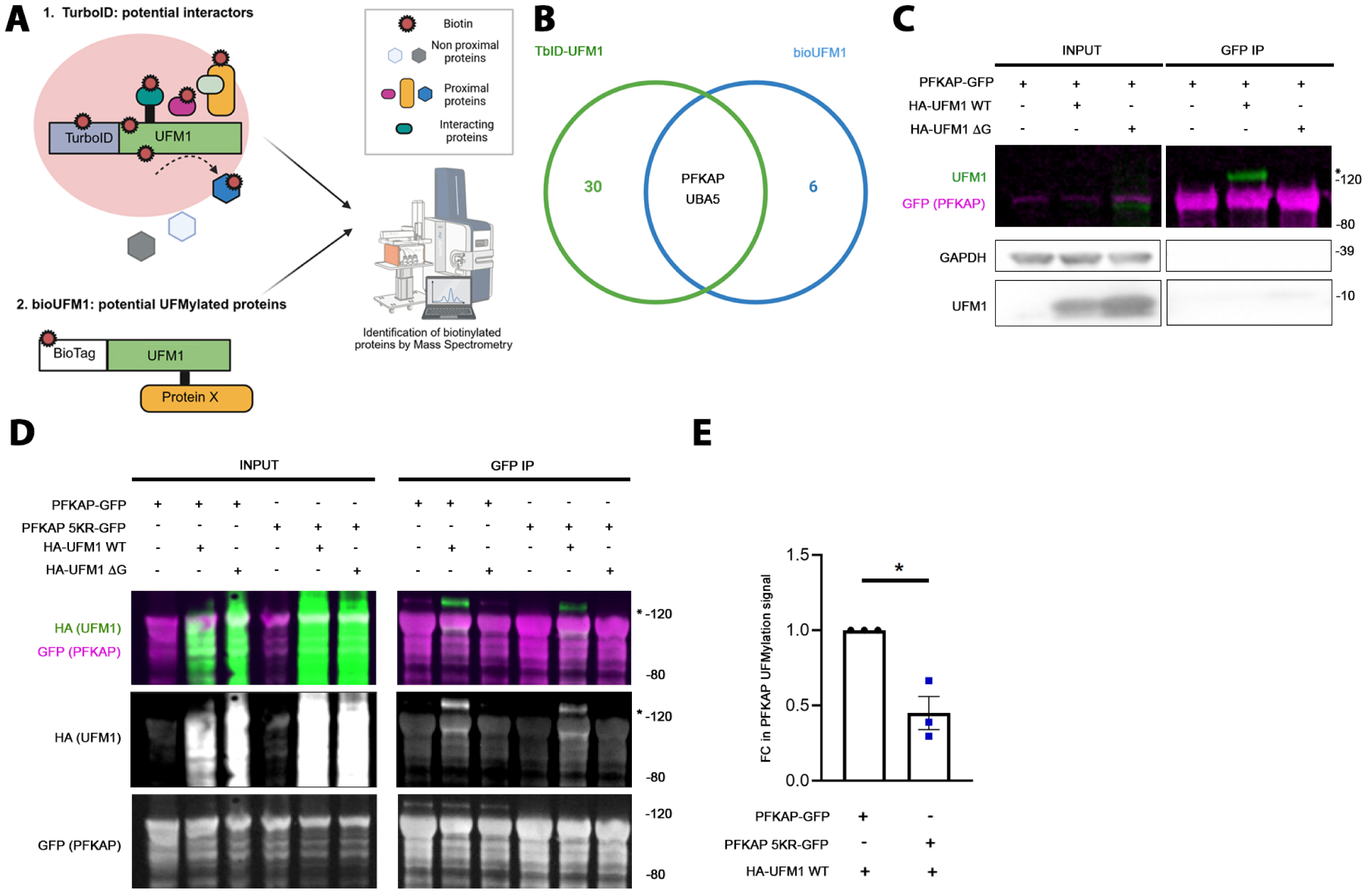
Identification of PFKAP as a target of UFMylation. **A.** Schematic overview of the TurboID-UFM1 (TbID-UFM1) and bioUFM1 techniques to identify UFMylated proteins in 22RV1 prostate cancer cells. In brief, proteins in the close proximity/potential interactors of UFM1 (TurboID) or proteins covalently modified by UFM1 (bioUFM1) are biotinylated and identified by Mass spectrometry (MS). *Created with BioRender.com*. **B.** Venn diagram showing the overlapping proteins identified both by the TbID-UFM1 and bio-UFM1 techniques. Thresholds: p value < 0.05 and fold change > 1 comparing TbID-UFM1 *vs.* TbID and bioUFM1 *vs.* BirA. **C.** Validation western blot analysis of exogenous PFKAP UFMylation. HEK293FT cells were transfected with PFKAP-GFP alone or together with functional UFM1 (HA-UFM1 WT) or conjugation-defective UFM1 (HA-UFM1 ΔG). PFKAP pulldown was carried out using GFP beads and membranes were probed with the indicated antibodies. UFMylated PFKAP is shown in green (UFM1) and unmodified PFKAP-GFP in purple (GFP). Molecular weight markers (kDa) are shown to the right. * marks the band of UFMylated PFKAP (in green). IP: immunoprecipitation. **D.** UFMylation analysis upon pulldown of wildtype (PFKAP WT-GFP) or lysine mutant PFKAP (K(281, 287, 395, 625, 736)-to-R; PFKAP 5KR-GFP) in HEK293FT cells expressing functional UFM1 (HA-UFM1 WT) or conjugation-defective UFM1 (HA-UFM1 ΔG). UFMylated PFKAP is shown in green (HA) and unmodified PFKAP-GFP in purple (GFP). Molecular weight markers (kDa) are shown to the right. * marks the bands of potential mono-UFMylated PFKAP. IP: immunoprecipitation. **E.** Quantification of the immunoreactivity of UFMylated exogenous (PFKAP WT-GFP) *vs.* PFKAP 5KR-GFP in the presence of HA-UFM1 WT in 3 independent western blot analyses (including the one shown in Fig. 3D). The graph represents the mean fold change in band intensity (*) of UFMylated exogenous PFKAP 5KR-GFP normalized to UFMylated PFKAP WT-GFP quantified by FiJi. P value was obtained by one-sample t-test.

**Fig. 4.**
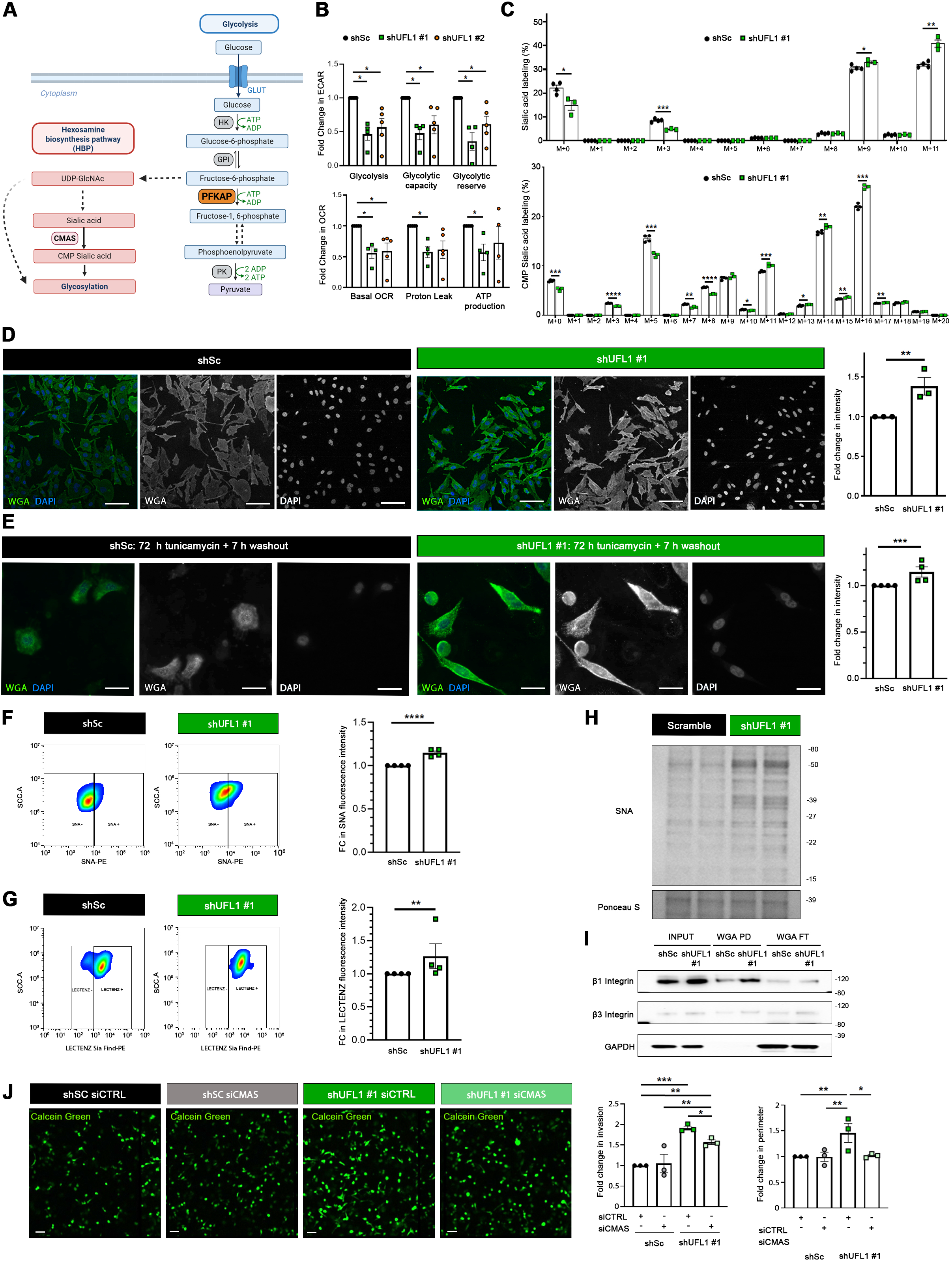
*UFL1* silencing increases hexosamine biosynthesis and protein glycosylation. **A.** Schematic representation of glycolysis and hexosamine biosynthesis pathways. *Created with BioRender.com*. **B.** Extracellular acidification rate (ECAR, top graph) and oxygen consumption rate (OCR, bottom graph) analysis of the Glycolysis Stress test in control (shSc) and *UFL1*-silenced (shUFL1 #1 and shUFL1 #2) PC3 cells by Seahorse analysis (n=5) related to shSc. P value was obtained by ordinary one-way ANOVA with multiple comparison analysis. **C.** Fractional contribution of ^13^C_6_-glucose to sialic acid (top) and CMP-sialic acid (bottom) carbon labelling upon 24h with ^13^C_6_-glucose, analyzed by LCMS in control (shSc) and *UFL1*-silenced (shUFL1 #1) PC3 cells (n=4 for shSc and n=3 for shUFL1 #1 cells). M+n represents the ^13^C labelled from glucose. P value was obtained by multiple unpaired t-tests. **D.** Representative micrographs and quantification of the levels of β-1,4-GlcNAc and sialic acid-linked residues in control (shSc) and *UFL1* silenced (shUFL1 #1) PC3 cells using wheat germ agglutinin (WGA) staining (n=4). Green, WGA (β-1,4-GlcNAc- and sialic acid-linked proteins); blue, DAPI nuclear staining. P value was obtained by one-sample t-test. Scale bar 15 μm. **E.** Representative micrographs and quantification of the levels of β-1,4-GlcNAc and sialic acid-linked residues in control (shSc) and *UFL1*-silenced (shUFL1 #1) PC3 cells after 72h of tunicamycin pretreatment (0.05 μg/ml) measured at 7h upon tunicamycin washout using wheat germ agglutinin (WGA) staining (n=4). Green, WGA (β-1,4-GlcNAc- and sialic acid-linked proteins); blue, DAPI nuclear staining. P value was obtained by one-sample t-test. Scale bar 30 μm. **F-G.** Representative flow cytometry histograms showing the expression of SNA (F) or Lectenz (G) in shUFL1 #1 cells compared to control cells (shSc) (left). On the right, graphs representing the fold change in SNA intensity (F) and Lectenz intensity (G) quantified by FlowJo software. P values were obtained by one-sample t-test. **H.** Representative western blot of two replicates of total lysates of control (shSc) and *UFL1*-silenced (shUFL1 #1) PC3 cells. Lectin SNA incubation and ponceau S staining as loading control were used. Molecular weight markers (kDa) are shown to the right. (n=5). **I.** Representative western blot showing the protein expression levels of glycosylated integrins β1 and β3 (elution) after WGA-mediated isolation of β-1,4-GlcNAc- and sialic acid-linked proteins from total lysates of control (shSc) and *UFL1*-silenced (shUFL1 #1) PC3 cells (n=5). Total levels of integrins β1 and β3 from the whole cell lysate and GAPDH as loading control are shown. PD: pulldown and FT: flowthrough. **J.** Representative images (left) and quantification (area and perimeter, right) of the invasive ability of control (shSc) and *UFL1*-silenced (shUFL1 #1) PC3 cells with functional (siCTRL) or inactive (siCMAS) hexosamine biosynthesis pathway in a collagen:matrigel 3D matrix. The total area occupied by calcein green-stained cells and their perimeter was measured using ImageJ (n=3). P value was obtained by repeated measures ANOVA with multiple comparison analysis.

Glycosylation of integrins is a critical process in dissemination of tumor cells^34^, a phenomenon that relies on the rewiring glucose metabolism towards hexosamine biosynthesis^25^. We thus monitored integrin glycosylation in our prostate cancer cells with dysfunctional UFMylation. We isolated all β-1,4-N-acetylglucosamine- and sialic-acid-linked proteins from PC3 cells using WGA pulldown and immunoblotted for integrin β1 and β3. Notably, we recovered more glycosylated integrin β1 and β3 in *UFL1*-silenced cells (**Fig. 4I**). These results indicate that impaired UFMylation could promote cancer cell invasion through increased hexosamine biosynthesis pathway and glycosylation of pro-invasive surface proteins, such as integrins.

To assess whether the hexosamine biosynthesis pathway contributes to the invasive phenotype of UFMylation deficient prostate cancer cells, we transiently silenced (using siRNA) N-acylneuraminate cytidylyltransferase (CMAS) in control (shSc) or *UFL1*-silenced (shUFL1) cells. Knockdown of *CMAS* tempered the increase in sialylated proteins (**Extended Data Fig. 4F-G**). Importantly, impairment of the hexosamine biosynthesis pathway prevented the increase in invasiveness elicited by *UFL1* silencing (**Fig. 4J**) without affecting cell number in the duration of the invasion assay (**Extended Data Fig. 4H**). These data support a model where the enhanced invasiveness of UFMylation-deficient cells is dependent on rewiring of glucose metabolism towards production of glycosylation intermediaries. Collectively, these data provide evidence that reduction in the UFMylation machinery in prostate cancer reduces the UFMylated moiety of a novel substrate, PFKAP, and redirects the flux of glucose towards hexosamine biosynthesis to support protein glycosylation, cell invasion and metastasis (**Extended Figure 5**).

## METHODS

### Cell cultures

Prostate cancer cell lines of human origin, PC3 (ACC465) and DU145 (ACC261) were purchased from and authenticated by Leibniz Institut DMSZ (Deutsche Sammlung von Mikroorganismen und Zellkulturen GmbH). Prostate cancer cell lines of human origin 22RV1, were purchased from American Type Culture Collection, ATCC (CRL-2505). Human embryonic kidney 293FT (HEK293FT) cells were generously provided by the laboratory of Dr. Rosa Barrio. PC3, DU145 and HEK293FT cells were cultured in Dulbecco’s modified eagle medium (DMEM) (Gibco, 41966-029). 22RV1 cells were maintained in Roswell park memorial institute (RPMI 1640, 11875-093) medium. Both culture mediums were supplemented with 10% inactivated foetal bovine serum (FBS) (Gibco, 10270-106) and 1% penicillin/ streptomycin (Gibco, 15140-122). Cell lines were tested for mycoplasma contamination routinely using MycoAlert detection Kit (Lonza, LT07-318). Cells were maintained at 37°C and 5% of CO_2_ pressure in a humidified chamber.

### Generation of stable cell lines

HEK293FT cells were used for lentiviral production. Lentiviral vectors expressing short hairpins (shRNAs) against human Scramble (shSc), UFM1(shUFM1 #1) and UFL1 (shUFL1 #1 and shUFL1 #2) were purchased from Sigma-Aldrich. The shRNA sequences are detailed in **Supplementary Table 7**. The infection was performed using standard procedures: HEK293FT cells were transfected with the appropriate lentiviral vectors using the calcium phosphate method and the viral supernatant plus protamine sulphate were used to infect PC3, DU145 or 22RV1 cells. Puromycin treatment (2 μg/ml; Sigma-Aldrich, P8833) was used for selection for 3 days. TurboID-UFM1 (TbID-UFM1) or TurboID control (TbID) lentiviral expression vectors were generated by replacing Cas9 in Lenti-Cas9-blast (Addgene #52962). For TurboID experiments, TbID-UFM1-P2A-blast or TbID-P2A-blast alone were transduced in 22RV1 cells and a stable population was selected with blasticidin (10 µg/ml) for 5 days (blasticidin was renewed after the first 3 days). For BioUFM1 experiments, bioUFM1-puro or BirA-puro plasmids were transduced in 22RV1 cells and a stable population was selected with puromycin (2 μg/ml; Sigma-Aldrich, P8833) for 3 days.

### Mice

All mouse experiments were carried out following the ethical guidelines established by the Biosafety and Animal Welfare Committee at CIC bioGUNE. The procedures employed were carried out following the recommendations from the Association for Assessment and Accreditation of Laboratory Animal Care (AAALAC). Mice were maintained in a controlled environment, with standard 12:12 light:dark cycles, 30-50% of humidity and controlled temperature at 22±2°C. Diet and water were provided *ad libitum*. At experimental endpoint, mice were sacrificed by CO2 inhalation followed by cervical dislocation. For intracardiac assays, 1.5 x 10^5^ PC3 GFP-Luc cells expressing shSc or shUFL1 #1 cells were injected intracardiacally into 8 Nu/Nu immunodeficient males of 6–12 weeks of age. Tumor growth and dissemination was followed by measuring bioluminescence for up to 25 days with IVIS technology (PerkinElmer). Intra-orbital injections of 50 μL luciferase at 15 mg/ml were used during the follow-up. Luciferase signal and tumor weight were measured. Cell dissemination was analyzed in mouse organs *ex vivo* by IVIS.

### Plasmid constructs and transient overexpression

All plasmid constructs are detailed in **Supplementary Table 7**. They were all verified by Sanger sequencing. Details of vector assembly are available on request. bioUFM1, bioUb-BirA, BirA, TbID-UFM1, TbID, HA-UFM1 WT, HA-UFM1 ΔG, HA-PFKAP, BirA-UBA5 (E1), BirA-UFC1 (E2), BirA-UFL1 (E3) and DDRGK1-YFP plasmids were generously provided by the laboratory of Dr. Rosa Barrio. pLVX neo PFKP-GFP^23^ was a gift from Gaudenz Danuser (Addgene plasmid # 116940 ; http://n2t.net/addgene:116940; RRID:Addgene_116940); pLVX neo PFKP-GFP K281R^23^ was a gift from Gaudenz Danuser (Addgene plasmid # 138288 ; http://n2t.net/addgene:138288 ; RRID:Addgene_138288). pLVX neo PFKP 5KR-GFP was generated by inserting PFKAP 5KR amplicon (purchased from Genscript) into Addgene plasmid # 116940. For transient plasmid overexpression, HEK293FT cells were transfected with the appropriate vectors using the calcium phosphate method. For transfection of siRNAs in different cancer cell lines, Lipofectamin RNAiMAX Transfection Reagent (Invitrogen, 13778150) was used, according to the manufacturer’s instructions. The siRNAs used in this study are described in **Supplementary Table 7.**

### Proliferation assays

Proliferation assays were performed by plating 5000 PC3 and 10000 DU145 and 22RV1 cells in triplicate in 12-well dishes and fixing them with 10% formalin (Avantor) after adhesion (day 0) and after having let them grow for 6 days. Cells were stained with crystal violet as previously described ^35^ and quantified after the resuspension of the crystals in 10% acetic acid by measuring the absorbance at 595nm.

### Foci growth assays

500 PC3 cells/well were seeded in a 6-well plate in triplicate. Cells were fixed with 10% formalin (Avantor) and stained with 0.1% crystal violet (Sigma-Aldrich) in 20% methanol after 10 days. Colonies were quantified using ImageJ^36^.

### Invasive growth assays

Cancer cells were tested for *in vitro* invasion either as loose cells or spheroids. For the invasion assay with loose cells^25^, 50,000 cells were embedded in a 50:50 mix of growth factor-reduced Matrigel (BD Biosciences) and collagen I (Life Technologies) and seeded onto 35 mm glass bottom culture dishes (MatTek). Cells were allowed to invade for 24h (DU145), 48 h (PC3) and 72h (22RV1), then they were stained with calcein green (Life Technologies) for 1h, washed with PBS and immediately imaged with SP8 confocal microscope (Leica). Images were acquired as three-dimensional scans with 10 μm Z-steps and processed with LAS X software (Leica) to obtain maximum projection images. Quantification of invasive area and invasive distance was performed on the maximum projection images using FiJi^37^. For relative invasive growth experiments using spheroids^38^, 100 PC3 cells were grown upside down in suspension with 20 µl 6% methylcellulose (Sigma-Aldrich, M0387) in DMEM drops. After 3 days, when the spheres were formed, they were embedded in a collagen-based medium containing 55% collagen I (Advanced BioMatrix PureCol, 5005), 20% DMEM 5X, 21.3% H_2_O_2_ and 3% 0.1 N NaOH, and photos were taken after collagen polymerization (0h time point) using an Olympus IX-83 inverted microscope operated by CellSens software. 5 experimental replicates were used for each condition. After 5 days, photos were taken, and the relative invasive growth was calculated as area difference on day 5 *vs.* day 0 using FiJi.

### Transwell assay

For transwell assay, 1 × 10^4^ PC3 cells were seeded in the upper chamber of a Corning BioCoat Matrigel Invasion Chamber (0.5 ml; inserts 6.4 mm, 8 μm pore size; Corning Costar, 354480) in serum-free medium. The lower chamber was loaded with 0.75 ml medium supplemented with 10% FBS. After 24h of incubation at 37°C with 5% CO_2_, the migrated cells in the membrane were fixed with formaline for 15 min, permeabilized with 0.2% TX-100 for 15 min, washed with PBS twice and stained with 0.5 µg/ml DAPI for 10 min. Images were obtained in an automated manner with the ScanR acquisition software controlling a motorized Olympus IX-83 wide-field microscope. Images from 3 independent experiments were then processed using the ScanR image analysis software.

### TurboID and bioUFM1 pulldowns

Using TurboID^21,39^ and bioUFM1^22^ methods, proteins in close proximity to UFM1 and potential UFM1 substrates, respectively, were biotinylated and isolated by streptavidin-bead pulldowns. 4 × 10^6^ 22RV1 cells were seeded in 5 × p150mm dishes in triplicates. Briefly, 24 hr after seeding, medium was supplemented with biotin at 50 µM. Cells were collected 2 h (for TurboID) and 24h (for bioUFM1) after biotin treatment, washed 3 times on ice with cold phosphate buffered saline (PBS) and scraped in lysis buffer [8 M urea, 1% SDS, 1x protease inhibitor cocktail (Roche, 05892970001), 60 mM NEM in 1x PBS; 2 ml per p150mm dish]. At room temperature, samples were sonicated and cleared by centrifugation. Extracted proteins were quantified using a Pierce BCA Protein Assay Kit (Thermo Scientific, 23225). Cell lysates were incubated overnight with 200 µl of equilibrated NeutrAvidin-agarose beads (Thermo Scientific). Beads were subjected to stringent washes using the following washing buffers (WB), all prepared in PBS: WB1 (8 M urea, 0.25% SDS); WB2 (6 M Guanidine-HCl); WB3 (6.4 M urea, 1 M NaCl, 0.2% SDS), WB4 (4 M urea, 1 M NaCl, 10% isopropanol, 10% ethanol and 0.2% SDS); WB5 (8 M urea, 1% SDS); and WB6 (2% SDS). For elution of biotinylated proteins, beads were heated at 99°C in 50 µl of Elution Buffer (4x Laemmli buffer, 100 mM DTT). Beads were separated by centrifugation (14000 rpm, 2min) through a Vivaclear mini 0.8 µm PES column (Sartorius, VK01P042). Samples were used for mass spectrometry (MS) analysis as well as for western blotting for validation of selected candidates.

### GFP immunoprecipitation

For the identification of PFKAP UFMylation sites by MS, 5 × 10^6^ HEK293FT were seeded in 15 × p150 mm dishes per condition. 4h after seeding, cells were transiently transfected with pLVX neo PFKP-GFP (Addgene; Plasmid #116940) and pRK5-HA-UFM1 (kindly donated by Dr. James D. Sutherland, CIC bioGUNE). For the rest of the GFP pull downs, 4 × 10^6^ HEK293FT were seeded in 1 × p100 mm dishes per condition. 4 h after seeding, cells were transiently transfected with the corresponding plasmids detailed in **Supplementary Table 7**. Cell culture media was refreshed after 24 h. 48 h after transfection, cells were washed two times with PBS, and lysed in 1.5 ml (for MS experiments) or 1ml (for the rest of experiments) of lysis buffer ((50 mM Tris-HCl [pH 7.65], 150 mM NaCl, 1 mM EDTA, 0.5% Triton X-100, 50 µM PR-619 DUB inhibitor and protease inhibitors (Roche, 05892970001) in water)). Lysates were spun down at 4200 rpm for 30 min for MS samples or at 16400 rpm for 15 min for the rest of experiments. Extracted proteins were quantified using a Pierce BCA Protein Assay Kit (Thermo Scientific, 23225). After 50 µl of supernatant (input) was saved, the rest was incubated 3h with 200 µl (for MS experiments) or 50 µl (for the rest of experiments) of pre-washed GFP-Trap resin (Chromotek, GTA-20) in a rotating wheel. Beads were washed once with dilution buffer ((10 mM Tris-HCl [pH 7.65], 150 mM NaCl, 1 mM EDTA, 50 µM PR-619 DUB inhibitor and protease inhibitors (Roche, 05892970001) in water)), twice with WB5 (8 M urea, 1% SDS in PBS) and once with 1% SDS in PBS. Beads were centrifuged at 2000 rpm for 2 min after each wash. For elution, samples were boiled for 5 min at 95°C in 50 µl elution buffer (4x Laemmli buffer, 100 mM DTT).

### Mass spectrometry proteomics

Analysis was done in 22RV1 cells stably expressing TbID-UFM1 or TbID and Bio-UFM1 or BirA. Three independent pulldown experiments (20 × 10^6^ cells per replicate) were analyzed by MS. Samples eluted from the NeutrAvidin beads were separated in SDS-PAGE (50% loaded) and stained with Sypro-Ruby (Biorad) according to manufacturer’s instructions. Entire gel lanes were excised, divided into pieces and in-gel digested with trypsin. Recovered peptides were desalted using stage-tip C18 microcolumns (Zip-tip, Millipore) and resuspended in 0.1% FA prior to MS analysis. Samples were analyzed in a hybrid trapped ion mobility spectrometry – quadrupole time of flight mass spectrometer (timsTOF Pro with PASEF, Bruker Daltonics) coupled online to a Evosep ONE liquid chromatograph (Evosep). Samples (200 ng) were loaded onto the chromatograph and resolved using either the 60 SPD (approx. 21 minute gradient) or the 30 SPD (approx. 44 min gradient) protocols. Protein identification and quantification was carried out using MaxQuant software^40^. Searches were carried out against a database consisting of human entries (Uniprot/Swissprot), with precursor and fragment tolerances of 20 ppm and 0.05 Da. For the bio-UFM1 and Turbo-ID experiments, only proteins identified with at least two peptides were considered for further analysis. Bio-UFM1 and TbID-UFM1 data were loaded onto Perseus platform^41^ and further processed (Log_2_ transformation, selection of proteins with at least two valid values in at least one condition, imputation). A t-test was applied to determine the statistical significance (p value < 0.05) of the differences detected. Proteins with a fold change in intensity higher than 1 and a p value lower than 0.05 between TbID-UFM1 and TbID or between bioUFM1 and BirA were considered for further analysis. For the identification of PFKAP UFMylation residues, a custom modification considering an increase in 156.09 Da on K was added to the PEAKS X+ software pipeline (Bioinformatics Solutions). UFMylation was considered as variable, together with the ubiquitination of lysine (K) and the oxidation of M (carbamidomethylation of C was kept fixed). All hits with a FDR < 1% were considered and UFMyalted hit spectra was manually inspected.

### HA Immunoprecipitation

4 × 10^6^ HEK293FT were seeded in 1 × p100 mm dishes per condition. 4 h after seeding, cells were transiently transfected with the corresponding plasmids detailed in **Supplementary Table 7**. Cell culture media was refreshed after 24 h. HEK293FT-transfected cells were collected after 48 h, washed 3 times with PBS, and lysed in 1 ml of lysis buffer ((20 mM Tris-HCl [pH 7.5], 150 mM NaCl, 1 mM EDTA, 1mM MgCl2 0.1% TX-100, 1mM NaF, Na Orthovanadate and β-GP mix and protease inhibitor mixture (Roche, 05892970001)). All steps were performed at 4°C. Lysates were spun down at 16400 rpm for 15 min. Extracted proteins were quantified using a Pierce BCA Protein Assay Kit (Thermo Scientific, 23225). After 50 µl of supernatant (Input) was saved, the rest was incubated overnight with 100 µl of pre-washed HA agarose beads (Thermo Scientific, 26182) in a rotating wheel. Beads were washed five times with IP washing buffer (0.05% TBS-T in water). Beads were centrifuged at 2000 rpm for 2 min after each wash. For elution, samples were boiled for 5 min at 95°C in 50 µl elution buffer (4x Laemmli buffer, 100 mM DTT).

### β-1,4-GlcNAc- and sialic acid-linked protein isolation

Glycoprotein Isolation Kit WGA (Thermo Scientific, 10290154) was used to isolate the glycoproteins rich in N-acetylglucosamine and sialic acid from whole cell lysates according to the manufacturer’s protocol. Briefly, cellular pellets were extracted twice: first with RIPA buffer [(50 mM Tris-HCl [pH 8], 150 mM NaCl, 1% Igepal (I8896, Sigma-Aldrich), 0.1% SDS, 0.5 sodium deoxycholate and protease inhibitor mixture (Roche, 05892970001)] and spun down at 16400 rpm for 10 minutes; then, the obtained pellets were extracted with RIPA buffer + 0.1 % TX-100 and spun down at 16400 rpm for 10 minutes. Both extractions were pooled together. Extracted proteins were quantified using a Pierce BCA Protein Assay Kit (Thermo Scientific, 23225). Samples containing 1 mg of protein per condition were processed through the WGA Lectin Resin column. After glycoprotein capture, the elution was collected for western blot probing.

### Protein extraction and western blot analysis

Cells were lysed in cold RIPA buffer supplemented with protease inhibitor cocktail (Roche, 05892970001). Lysates were kept on ice for 15 min, vortexing every 5 min and then, cleared by centrifugation (16400 rpm for 10 min at 4°C). Supernatants were collected, and protein contents were quantified by a Pierce BCA Protein Assay Kit (Thermo Scientific, 23225). Protein extraction western blot was performed as previously described^42^. Briefly, samples were run in 4-12% gradient NuPage (Life Technologies, WG1403BX10) or 4-20% gradient Tris-Glycine Plus (Life Technologies, WXP42020BOX) precast gels in MOPS buffer. After transfer to nitrocellulose, membranes were blocked in 5% milk in TBS and 0.1% Tween-20 or Carbo-Free Blocking Solution (10x Concentrate, SP-5040-125) at 1X in TBS and 0.1% Tween-20 for SNA western blot probing. Primary antibodies were incubated overnight at 4°C, and secondary antibodies were incubated for 1 h at room temperature. Primary antibodies for UFL1 (Sigma-Aldrich, HPA030559), UFM1 (Abcam, ab109305), GAPDH (Cell Signaling, 2118L), HA (Cell Signalling Technologies, 3724), GFP (Roche, 11814460001), PFKAP (Proteintech, 13389-1-AP), BirA (Sino Biological, 11582-T16-100), β1-integrin (Cell Signaling, 34971S) and β3-integrin (Cell Signaling Technology, 13166T) were used at a 1:1000 dilution. Biotinylated SNA (Vectorlabs, B-1305-2) was also used at 1:1000 dilution. Secondary HRP-conjugated anti-rabbit or anti mouse antibodies (Jackson ImmunoResearch, 111-035-144), biotin-HRP (Cell Signaling Technology, 7075), Streptavidin-HRP (Thermo Scientific, N100) or Alexa 488 and 647-conjugated mouse antibody were used at a 1:5000 dilution.

### N-glycosylation immunofluorescence

For non-treated cells, PC3 shSc and shUFL1 #1 cells were seeded at 5 × 10^5^ cells per well on a µ-Slide 8 Well (Ibidi, 80826). Cells were then fixed in paraformaldehyde 4% in PBS for 20 min at RT. Cells were incubated with wheat germ agglutinin (WGA) Alexa Fluor 555 (Thermo Scientific, W32464) at final concentration of 5 μg/ml in Hank’s balanced salt solution (HBSS) without phenol red. Samples were washed three times with PBS and incubated with 1 μg/ml of Hoechst (Life Technologies, H3570) for 5 min at RT. For tunicamycin treated cells, PC3 shSc and shUFL1 #1 cells were treated for 72 h with 0.05 ug/ml tunicamycin (Merk Life Science, T7765). Tunicamycin was washed out and the cells still attached to the flask were harvested and seeded at 2×10^5^ on 11 mm round glass coverslip previously coated with 10% Poly-L-lysine in PBS (Sigma-Aldrich, P8920) and placed in 24-well plates for 7h. Cells adhering to chambers or coverslips were incubated for 10 min at 37°C with wheat germ agglutinin (WGA) Alexa Fluor 555 (Thermo Scientific, W32464) at final concentration of 5 μg/ml in Hank’s balanced salt solution (HBSS) without phenol red. Samples were then fixed in paraformaldehyde 4% in PBS for 20 min at RT. At last, samples were washed three times with PBS and incubated with 1 μg/ml of Hoechst (Life Technologies, H3570) for 5 min at RT. Imaging was performed in an automated manner with the ScanR acquisition software controlling a motorized Olympus IX-83 wide-field microscope. Images from 3 independent experiments were then processed using the ScanR image analysis software.

### Seahorse Glycolysis Stress metabolism measurements

5×10^3^ PC3 and 22RV1, and 10×10^3^ DU145 shSc, shUFL1 #1 and shUFL1 #2 cells were plated in 24-well Seahorse XF cell culture plates (Seahorse Bioscience, 100882-004). Seahorse XFe24 sensor cartridge plates were hydrated with the XF Calibrant (Seahorse Bioscience) the day before the analysis and incubated overnight at 37 °C without CO2. Before the bioenergetics measurements, cells were washed and incubated for 1 h with Glycolysis Stress media. Glycolysis Stress media contained XF Base Medium (minimal DMEM, Seahorse Bioscience, 103575-100) supplemented with 2 mM L-glutamine (Seahorse Bioscience, 103579-100). The extracellular acidification rate (ECAR), representative of the glycolytic capacity, and the oxygen consumption rate (OCR), representative of the mitochondrial respiration, were determined using the XFe Extracellular Flux Analyzer (Agilent/Seahorse Bioscience). The glycolytic metabolism was determined by the sequential injection of 10 mM D-(+)-glucose (Sigma-Aldrich, G8644), 1.5 µM of the ATP synthase inhibitor oligomycin (Sigma-Aldrich, 75351) to inhibit mitochondrial respiration and force the cells to maximize their glycolytic capacity, and 50 mM 2-deoxy-D-glucose (2-DG) (Sigma-Aldrich, D8375), a competitive inhibitor of the first step of glycolysis. The concentrations indicated for each injection represent the final concentrations in the wells. At least three measurement cycles (3 min of mixing + 3 min of measuring) were completed before and after each injection. The OCR and ECAR were calculated using Wave software v2.6.3 (Agilent/Seahorse Bioscience). Following the manufacturer’s instructions, energy metabolism was normalized according to cell number using crystal violet staining.

### Liquid chromatography-mass spectrometry (LC-MS) for metabolite measurements

PC3 shSc and shUFL1 #1 cells (3×10^5^ per biological replicate) were treated for 24h with 25mM ^13^C_6_-labelled-D-glucose (Sigma-Aldrich, 389374). All labeling experiments were performed in media with 10% dialyzed serum. Metabolites for the subsequent mass spectrometry analysis were prepared by quenching the cells in liquid nitrogen followed by a cold two-phase methanol-water-chloroform extraction^43^. Phase separation was achieved by centrifugation at 4°C. The methanol-water phase containing polar metabolites was separated and dried using a vacuum concentrator at 4°C. Dried metabolite samples were stored at −80°C until analysis. The protein interphase was also dried down and dissolved in 200 μl 0.2 M potassium hydroxide, and the protein concentration was then quantified using the Pierce BCA Protein Assay Kit (Thermo Scientific, 23225). For the detection of metabolites by LC-MS, a Dionex UltiMate 3000 LC System (Thermo Scientific) with a thermal autosampler set at 4°C, coupled to a Q-Exactive Orbitrap mass spectrometer (Thermo Fisher Scientific) was used. Samples were resuspended in 50 μL of water and a volume of 10 μl of sample was injected into a C18 column (Acquity UPLC HSS T3 1.8 μm 2.1×100 mm). The separation of metabolites was achieved at 40 °C with a flow rate of 0.25 ml/min. A gradient was applied for 40 min (solvent A: 10mM Tributyl-Amine, 15 mM acetic acid – solvent B: Methanol) to separate the targeted metabolites (0 min: 0% B, 2 min: 0% B, 7 min: 37% B, 14 min: 41% B, 26 min: 100% B, 30 min: 100% B, 31 min: 0% B; 40 min: 0% B. The MS operated in negative full scan mode (m/z range: 70–1050 and 300–700 from 5 to 25 min) using a spray voltage of 4.9 kV, capillary temperature of 320 °C, sheath gas at 50.0, auxiliary gas at 10.0. Data was collected using the Xcalibur software (Thermo Scientific) and analyzed with Matlab for the correction of protein content and natural abundance, but also to determine the isotopomer distribution using the method developed by^44^. Metabolites abundances were corrected by protein content.

### Flow cytometry for lectin binding

5 × 10^5^ PC3 shSc and shUFL1 #1 PC3 cells were seeded per well of a 96-well round-bottomed microtiter plates per condition. Pelleted cells (450 x g, 5 min) were washed three times with 1% BSA in PBS. Washed cells were incubated with biotinylated SNA (Vectorlabs, B-1305-2) or SiaFind α2,3-Specific Lectenz (Lectenz Bio, SP2302B) at 5 µg/mL in 100 µL per well of 1% BSA in PBS for 1 h at 4 °C. After three washes at 450 x g for 5 min, cells were incubated in 100 µL containing Streptavidin PE (1:200) (BD Biosciences, 554061) for 30 min at 4 °C. Cells were then washed and resuspended in 200 µL of 1 % BSA in PBS with 0.5 µg/ml DAPI before acquisition by FACSAttune (Thermo Fisher Scientific). Results were analyzed using FlowJo (BD Biosciences).

### qRT-PCR

RNA was automatically extracted using Maxwell RSC instrument (Promega) according to manufactureŕs instructions. 1 µg of the extracted RNA was used for complementary DNA (cDNA) synthesis using Maxima™ H Minus cDNA Synthesis Master Mix (Invitrogen, M1682). QS6 (Life Technologies) system was used for RT-qPCR analysis. Applied biosystems TaqMan probes: *CMAS* (Hs00218814_m1) and *GAPDH* (Hs02758991_g1) were used.

### Bioinformatic analysis

Gene differential expression analysis was performed using the *limma*^45^ package across five publicly available prostate cancer datasets present in Cancertool^46^: Grasso, Lapointe, Taylor, Tomlins, and Varambally. For each dataset, a linear model was fitted using a design matrix without an intercept to model expression differences between sample groups. Log_2_ fold changes were estimated by specifying contrasts of interest (e.g., “Primary tumor-Metastasis”), followed by empirical Bayes moderation of the standard errors using the eBayes function. Gene ontology analysis from patient data sets was obtained using the web-based interface DAVID^47,48^. Cancer genomics data sets were analyzed using cbioportal^49^.

### Statistical analysis

Sample size was not predetermined using any statistical method and experiments were not randomized. Investigators were not blinded during experiments or outcome assessment. All the experiments were performed with at least three biological replicates. N values represent the number of independent biological experiments or the number of individual mice. In *in vitro* experiments, one-sample t-test was applied for one-component comparisons with control with a hypothetical value of 1 and unpaired or paired t-test for two-component comparisons. For *in vivo* experiments, Mann-Whitney test was used. For more than two-component comparisons, ordinary or repeated measures and two-way ANOVA with multiple comparison analysis was performed. For contingency studies, Chi square analysis was applied. Unless specified otherwise, two-tailed statistical analysis was applied for experimental design without predicted result. When suitable, one-tailed statistical analysis was applied for validation or hypothesis-driven experiments. The results are presented as the mean ± standard error of the mean (SEM). The confidence level used for all statistical analysis was 95% (p value = 0.05). p value: * p < 0.05, ** p < 0.01, *** p < 0.001. Statistical data analysis was performed using GraphPad Prism version 8.2 (GraphPad Software). Details on sample size, statistical tests and post-tests are presented in the figure legends.

## SEPARATE DATA STATEMENT

All data supporting the findings of this study are available within article and the Supplementary Information, and from the corresponding author on reasonable request.

## CODE AVAILABILITY STATEMENT

All the custom code used in the study is available from the corresponding author on reasonable request.

## ACKNOWLEDGEMENTS

We are grateful to the Carracedo lab for valuable input. Laura Bozal-Basterra was supported by the AECC Foundation (POSTD19048BOZA), the Boehringer Ingelheim Fonds Travel Grant, the EACR Travel Fellowship (application #784) and the EMBO Scientific Exchange Grant (#10477). This study was developed in the framework of the AECC Excelencia PREMETACAN funding scheme. The work in the Carracedo lab is supported by the Basque Department of Industry, Tourism and Trade (Elkartek), the MICINN (PID2022-141553OB-I0 (FEDER/EU); Fundación Cris Contra el Cáncer (PR_EX_2021-22), Severo Ochoa Excellence Accreditation (CEX2021-001136-S), iDIFFER network of Excellence (RED2022-134792-T), the AstraZeneca Foundation (award to young investigators 2023), Leonardo Grant for Scientific Research and Cultural Creation (Beca Leonardo LEO23-2-10984-BBM-TRA-261), and the European Research Council (Consolidator Grant 819242). CIBERONC was co-funded with FEDER funds and funded by ISCIII. I.M. is supported by Fundación Cris Contra El Cáncer (PR_TPD_2020-19) and a Ramón y Cajal contract (RYC2023-044682-I) funded by the MCIN. P.A.M. was supported by a Marie Sklodowska-Curie Actions individual fellowship (MSCA-IF-2018-839896) and has received funding from European Union (ERC-StG-101116912) and Beug Foundation. S.-M.F. acknowledges funding from, FWO Projects, Beug Foundation, Fonds Baillet Latour, KU Leuven, Stichting tegen Kanker and Interuniversity BOF (iBOF) program. M.G.V.H. acknowledges support from the Ludwig Center at MIT, the MIT Center for Precision Cancer Medicine, and the NCI (R35CA242379, R01CA259253, P30CA1405141). RB and JDS acknowledge PID2023-147399NB-I00 (MCIN/ AEI /10.13039/501100011033) and the Severo Ochoa Excellence Accreditation (CEX2021-001136-S). Illustrations in figures 1A, 3A, 3B, 4A Extended data Fig. 1D and Extended data Fig. 5 were created with BioRender.

## AUTHOR CONTRIBUTIONS

LB-B supervised, designed, performed and coordinated most *in vitro* and *in vivo* experiments (unless specified otherwise), analysed the data and wrote the manuscript. MC-S and AM-V contributed to the cell culture of different cell lines and *in vitro* experiments. AMF-C (supervised by SMF) and DRS (supervised by MVH) contributed to the design, performed and/or coordinated metabolomics analyses. MD, PA-M, OC, IA, AZ-L, AT and JF-G provided technical support and critical discussions. SG-L, MP-R, IM and NM-M provided bioinformatics support. KS, JE-O and JJ-B contributed to the design and provided technical support and scientific resources for the glycosylation assays. MA and II performed the proteomics experiments and/or analysed data (supervised by FE). SM-F and MVH contributed to critical scientific discussions. RB and JDS provided technical and intellectual support with the design and execution of biotin-based proteomics, and contributed to scientific discussions. AC designed the research strategy, directed the project, supervised the study and wrote the manuscript.

## COMPETING INTEREST DECLARATION

MVH discloses that he is a scientific advisor for Agios Pharmaceuticals, iTeos Therapeutics, Sage Therapeutics, Pretzel Therapeutics, Lime Therapeutics, Faeth Therapeutics, Droia Ventures, MPM Capital and Auron Therapeutics.

## CORRESPONDING AUTHOR LINE

Correspondence and requests for materials should be addressed to Arkaitz Carracedo.

## SUPPLEMENTARY INFORMATION

Supplementary Information is available for this paper:

**Supplementary Figure 1.** Identification of K281 and K287 lysine residues of PFKAP modified by UFM1 through mass spectrometry: GFP was immunoprecipitated from cell lysates of HEK293FT cells expressing PFKAP-GFP and HA-UFM1 WT, followed by mass spectrometry analysis.

**Supplementary Figure 2.** Identification of K395 lysine residue of PFKAP modified by UFM1 through mass spectrometry: GFP was immunoprecipitated from cell lysates of HEK293FT cells expressing PFKAP-GFP and HA-UFM1 WT, followed by mass spectrometry analysis.

**Supplementary Figure 3.** Identification of K625 lysine residue of PFKAP modified by UFM1 through mass spectrometry: GFP was immunoprecipitated from cell lysates of HEK293FT cells expressing PFKAP-GFP and HA-UFM1 WT, followed by mass spectrometry analysis.

**Supplementary Figure 4.** Identification of K736 lysine residue of PFKAP modified by UFM1 through mass spectrometry: GFP was immunoprecipitated from cell lysates of HEK293FT cells expressing PFKAP-GFP and HA-UFM1 WT, followed by mass spectrometry analysis.

**Supplementary Table 1.** This table is complementary and related to Fig.1A and Extended Data Fig.1A. It contains the raw and filtered transcriptomic data from four different prostate cancer cohorts.

**Supplementary Table 2.** This table is complementary and related to Fig.1B. It contains the results of the Gene Ontology analysis performed at *cancertool.org* for the 114 downregulated genes in metastasis.

**Supplementary Table 3.** This table is complementary and related to Fig. 3A-B and Extended Data Fig. 3A. It contains the mass spectrometry results of the TurboID-UFM1 (TbID-UFM1) technique in 22RV1 prostate cancer cells.

**Supplementary Table 4.** This table is complementary and related to Fig. 3A-B and Extended Data Fig. 3B. It contains the mass spectrometry results of the bioUFM1 technique in 22RV1 prostate cancer cells.

**Supplementary Table 5.** This table is complementary and related to Fig. 3D-E and Extended Data Fig. 3F. It contains the mass spectrometry results for the identification of the 5 lysine (K) residues of PFKAP modified by UFM1.

**Supplementary Table 6.** This table is complementary and related to Fig. 4C and Extended Data Fig. 4E. It contains information on sialic acid, CMP-sialic acid and UDP-GlcNac metabolites analysed in PC3 cultured cells by LCMS analysis.

**Supplementary Table 7.** This table is complementary and related to methods and contains information about the plasmids, shRNAs and siRNAs used in the manuscript.

## MAIN REFERENCES

1. Faubert, B., Solmonson, A. & DeBerardinis, R. J. Metabolic reprogramming and cancer progression. Science (1979) 368, (2020).

2. Komatsu, M., Chiba, T., Tatsumi, K., Iemura, S., Tanida, I., Okazaki, N., et al. A novel protein-conjugating system for Ufm1, a ubiquitin-fold modifier. EMBO J 23, 1977–1986 (2004).

3. Kang, S. H., Kim, G. R., Seong, M., Baek, S. H., Seol, J. H., Bang, O. S., et al. Two novel ubiquitin-fold modifier 1 (Ufm1)-specific proteases, UfSP1 and UfSP2. J Biol Chem 282, 5256–5262 (2007).

4. Millrine, D., Cummings, T., Matthews, S. P., Peter, J. J., Magnussen, H. M., Lange, S. M., et al. Human UFSP1 is an active protease that regulates UFM1 maturation and UFMylation. Cell Rep 40, 111168 (2022).

5. Wei, Y. & Xu, X. UFMylation: A Unique & Fashionable Modification for Life. Genomics Proteomics Bioinformatics 14, 140–146 (2016).

6. Yoo, H. M., Kang, S. H., Kim, J. Y., Lee, J. E., Seong, M. W., Lee, S. W., et al. Modification of ASC1 by UFM1 is crucial for ERα transactivation and breast cancer development. Mol Cell 56, 261–274 (2014).

7. Gong, Y., Wang, Z., Zong, W., Shi, R., Sun, W., Wang, S., et al. PARP1 UFMylation ensures the stability of stalled replication forks. Proc Natl Acad Sci U S A 121, e2322520121 (2024).

8. Qin, B., Yu, J., Nowsheen, S., Wang, M., Tu, X., Liu, T., et al. UFL1 promotes histone H4 ufmylation and ATM activation. Nat Commun 10, 1242 (2019).

9. Wang, Z., Gong, Y., Peng, B., Shi, R., Fan, D., Zhao, H., et al. MRE11 UFMylation promotes ATM activation. Nucleic Acids Res 47, 4124–4135 (2019).

10. Walczak, C. P., Leto, D. E., Zhang, L., Riepe, C., Muller, R. Y., DaRosa, P. A., et al. Ribosomal protein RPL26 is the principal target of UFMylation. Proc Natl Acad Sci U S A 116, 1299–1308 (2019).

11. Xu, A. & Barna, M. Cleaning up stalled ribosome-translocon complexes with ufmylation. Cell Res 30, 1–2 (2020).

12. Tatsumi, K., Yamamoto-Mukai, H., Shimizu, R., Waguri, S., Sou, Y.-S., Sakamoto, A., et al. The Ufm1-activating enzyme Uba5 is indispensable for erythroid differentiation in mice. Nat Commun 2, 181 (2011).

13. Yang, R., Wang, H., Kang, B., Chen, B., Shi, Y., Yang, S., et al. CDK5RAP3, a UFL1 substrate adaptor, is crucial for liver development. Development 146, (2019).

14. Zhou, J., Ma, X., He, X., Chen, B., Yuan, J., Jin, Z., et al. Dysregulation of PD-L1 by UFMylation imparts tumor immune evasion and identified as a potential therapeutic target. Proc Natl Acad Sci U S A 120, e2215732120 (2023).

15. Liu, J., Guan, D., Dong, M., Yang, J., Wei, H., Liang, Q., et al. UFMylation maintains tumour suppressor p53 stability by antagonizing its ubiquitination. Nat Cell Biol 22, 1056–1063 (2020).

16. Zhou, J., Ma, X., Xu, L., Liang, Q., Mao, J., Liu, J., et al. Genomic profiling of the UFMylation family genes identifies UFSP2 as a potential tumour suppressor in colon cancer. Clinical and translational medicine vol. 11 e642 Preprint at 10.1002/ctm2.642 (2021).

17. Franceschi, R., Iascone, M., Maitz, S., Marchetti, D., Mariani, M., Selicorni, A., et al. A missense mutation in DDRGK1 gene associated to Shohat-type spondyloepimetaphyseal dysplasia: Two case reports and a review of literature. Am J Med Genet A 188, 2434–2437 (2022).

18. Nahorski, M. S., Maddirevula, S., Ishimura, R., Alsahli, S., Brady, A. F., Begemann, A., et al. Biallelic UFM1 and UFC1 mutations expand the essential role of ufmylation in brain development. Brain 141, 1934–1945 (2018).

19. Wang, X., Xu, X. & Wang, Z. The Post-Translational Role of UFMylation in Physiology and Disease. Cells 12, (2023).

20. Gerakis, Y., Quintero, M., Li, H. & Hetz, C. The UFMylation System in Proteostasis and Beyond. Trends Cell Biol 29, 974–986 (2019).

21. Branon, T. C., Bosch, J. A., Sanchez, A. D., Udeshi, N. D., Svinkina, T., Carr, S. A., et al. Efficient proximity labeling in living cells and organisms with TurboID. Nat Biotechnol 36, 880–887 (2018).

22. Pirone, L., Xolalpa, W., Sigurðsson, J. O., Ramirez, J., Pérez, C., González, M., et al. A comprehensive platform for the analysis of ubiquitin-like protein modifications using in vivo biotinylation. Sci Rep 7, 40756 (2017).

23. Park, J. S., Burckhardt, C. J., Lazcano, R., Solis, L. M., Isogai, T., Li, L., et al. Mechanical regulation of glycolysis via cytoskeleton architecture. Nature 578, 621–626 (2020).

24. Akella, N. M., Ciraku, L. & Reginato, M. J. Fueling the fire: Emerging role of the hexosamine biosynthetic pathway in cancer. BMC Biol 17, 1–14 (2019).

25. Rossi, M., Altea-Manzano, P., Demicco, M., Doglioni, G., Bornes, L., Fukano, M., et al. Author Correction: PHGDH heterogeneity potentiates cancer cell dissemination and metastasis. Nature vol. 609 E8 Preprint at 10.1038/s41586-022-05226-7 (2022).

26. Buescher, J. M., Antoniewicz, M. R., Boros, L. G., Burgess, S. C., Brunengraber, H., Clish, C. B., et al. A roadmap for interpreting (13)C metabolite labeling patterns from cells. Curr Opin Biotechnol 34, 189–201 (2015).

27. Itkonen, H. M., Engedal, N., Babaie, E., Luhr, M., Guldvik, I. J., Minner, S., et al. UAP1 is overexpressed in prostate cancer and is protective against inhibitors of N-linked glycosylation. Oncogene 34, 3744–3750 (2015).

28. Munkley, J., Vodak, D., Livermore, K. E., James, K., Wilson, B. T., Knight, B., et al. Glycosylation is an Androgen-Regulated Process Essential for Prostate Cancer Cell Viability. EBioMedicine 8, 103–116 (2016).

29. Itkonen, H. M., Minner, S., Guldvik, I. J., Sandmann, M. J., Tsourlakis, M. C., Berge, V., et al. O-GlcNAc transferase integrates metabolic pathways to regulate the stability of c-MYC in human prostate cancer cells. Cancer Res 73, 5277–5287 (2013).

30. Albitar, M., Ma, W., Lund, L., Albitar, F., Diep, K., Fritsche, H. A., et al. Predicting Prostate Biopsy Results Using a Panel of Plasma and Urine Biomarkers Combined in a Scoring System. J Cancer 7, 297–303 (2016).

31. Elbein, A. D. The Use of Glycosylation Inhibitors to Study Glycoconjugate Function. Cell Surface and Extracellular Glycoconjugates 119–180 Preprint at 10.1016/B978-0-12-589630-6.50009-5 (1993).

32. Yoo, J., Mashalidis, E. H., Kuk, A. C. Y., Yamamoto, K., Kaeser, B., Ichikawa, S., et al. GlcNAc-1-P-transferase-tunicamycin complex structure reveals basis for inhibition of N-glycosylation. Nat Struct Mol Biol 25, 217–224 (2018).

33. Surani, M. A. Glycoprotein synthesis and inhibition of glycosylation by tunicamycin in preimplantation mouse embryos: compaction and trophoblast adhesion. Cell 18, 217–227 (1979).

34. He, M., Zhou, X. & Wang, X. Glycosylation: mechanisms, biological functions and clinical implications. Signal Transduct Target Ther 9, 194 (2024).

35. Torrano, V., Valcarcel-Jimenez, L., Cortazar, A. R., Liu, X., Urosevic, J., Castillo-Martin, M., et al. The metabolic co-regulator PGC1α suppresses prostate cancer metastasis. Nat Cell Biol 18, 645–656 (2016).

36. Schneider, C. A., Rasband, W. S. & Eliceiri, K. W. NIH Image to ImageJ: 25 years of image analysis. Nat Methods 9, 671–675 (2012).

37. Schindelin, J., Arganda-Carreras, I., Frise, E., Kaynig, V., Longair, M., Pietzsch, T., et al. Fiji: an open-source platform for biological-image analysis. Nat Methods 9, 676–682 (2012).

38. Crosas-Molist, E., Bertran, E., Rodriguez-Hernandez, I., Herraiz, C., Cantelli, G., Fabra, A., et al. The NADPH oxidase NOX4 represses epithelial to amoeboid transition and efficient tumour dissemination. Oncogene 36, 3002–3014 (2017).

39. Roux, K. J., Kim, D. I., Raida, M. & Burke, B. A promiscuous biotin ligase fusion protein identifies proximal and interacting proteins in mammalian cells. J Cell Biol 196, 801–810 (2012).

40. Tyanova, S., Temu, T. & Cox, J. The MaxQuant computational platform for mass spectrometry-based shotgun proteomics. Nat Protoc 11, 2301–2319 (2016).

41. Tyanova, S., Temu, T., Sinitcyn, P., Carlson, A., Hein, M. Y., Geiger, T., et al. The Perseus computational platform for comprehensive analysis of (prote)omics data. Nat Methods 13, 731–740 (2016).

42. Zabala-Letona, A., Arruabarrena-Aristorena, A., Martin-Martin, N., Fernandez-Ruiz, S., Sutherland, J. D., Clasquin, M., et al. mTORC1-dependent AMD1 regulation sustains polyamine metabolism in prostate cancer. Nature 547, 109–113 (2017).

43. van Gorsel, M., Elia, I. & Fendt, S.-M. (13)C Tracer Analysis and Metabolomics in 3D Cultured Cancer Cells. Methods Mol Biol 1862, 53–66 (2019).

44. Fernandez, C. A., Des Rosiers, C., Previs, S. F., David, F. & Brunengraber, H. Correction of 13C mass isotopomer distributions for natural stable isotope abundance. J Mass Spectrom 31, 255–262 (1996).

45. Ritchie, M. E., Phipson, B., Wu, D., Hu, Y., Law, C. W., Shi, W., et al. limma powers differential expression analyses for RNA-sequencing and microarray studies. Nucleic Acids Res 43, e47 (2015).

46. Cortazar, A. R., Torrano, V., Martin-Martin, N., Caro-Maldonado, A., Camacho, L., Hermanova, I., et al. CANCERTOOL: A Visualization and Representation Interface to Exploit Cancer Datasets. Cancer Res 78, 6320–6328 (2018).

47. Sherman, B. T., Hao, M., Qiu, J., Jiao, X., Baseler, M. W., Lane, H. C., et al. DAVID: a web server for functional enrichment analysis and functional annotation of gene lists (2021 update). Nucleic Acids Res 50, W216–W221 (2022).

48. Huang, D. W., Sherman, B. T. & Lempicki, R. A. Systematic and integrative analysis of large gene lists using DAVID bioinformatics resources. Nat Protoc 4, 44–57 (2009).

49. Cerami, E., Gao, J., Dogrusoz, U., Gross, B. E., Sumer, S. O., Aksoy, B. A., et al. The cBio cancer genomics portal: an open platform for exploring multidimensional cancer genomics data. Cancer Discov 2, 401–404 (2012).

