## Extended Data for "Loss of UFMylation supports prostate cancer metastasis and rewires cell metabolism towards hexosamine biosynthesis"

### Extended Data Figure 1

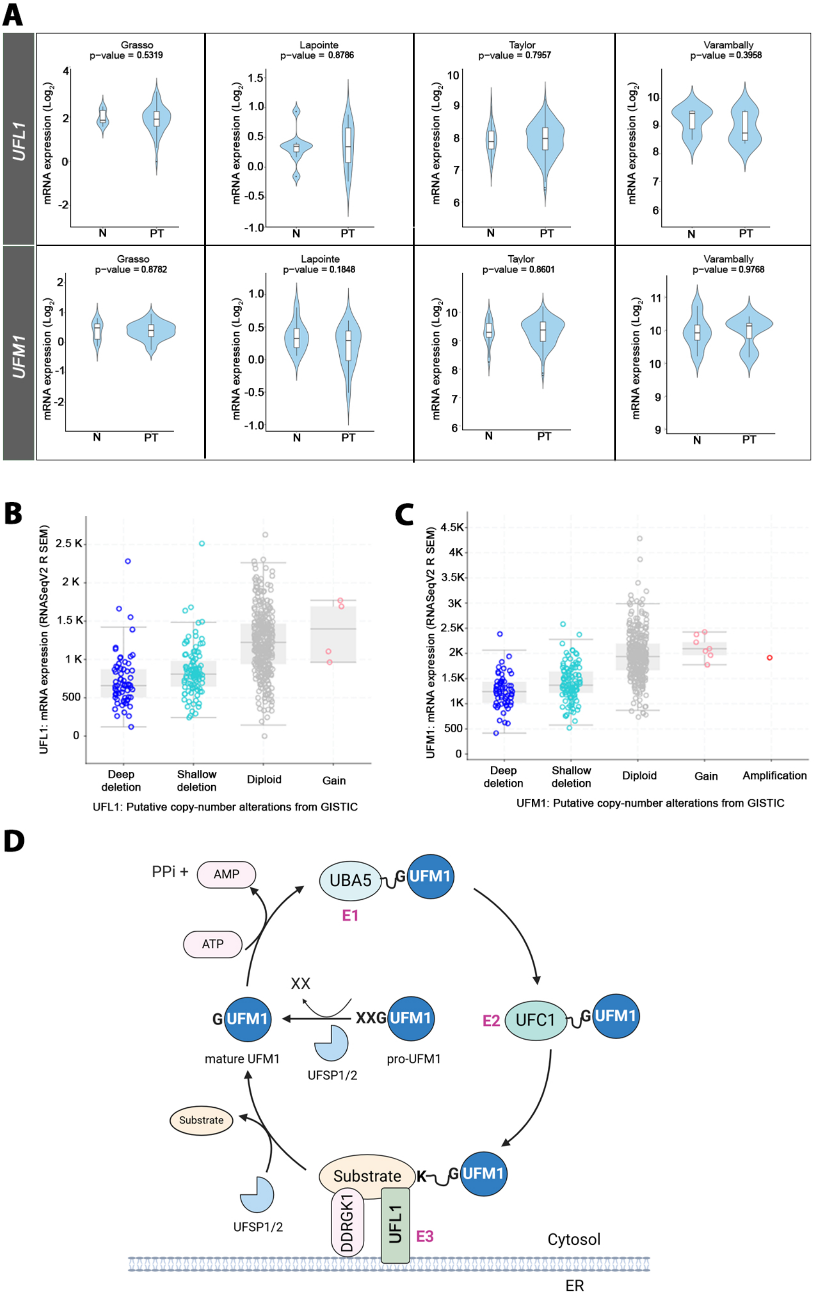

### Extended Data Figure 2

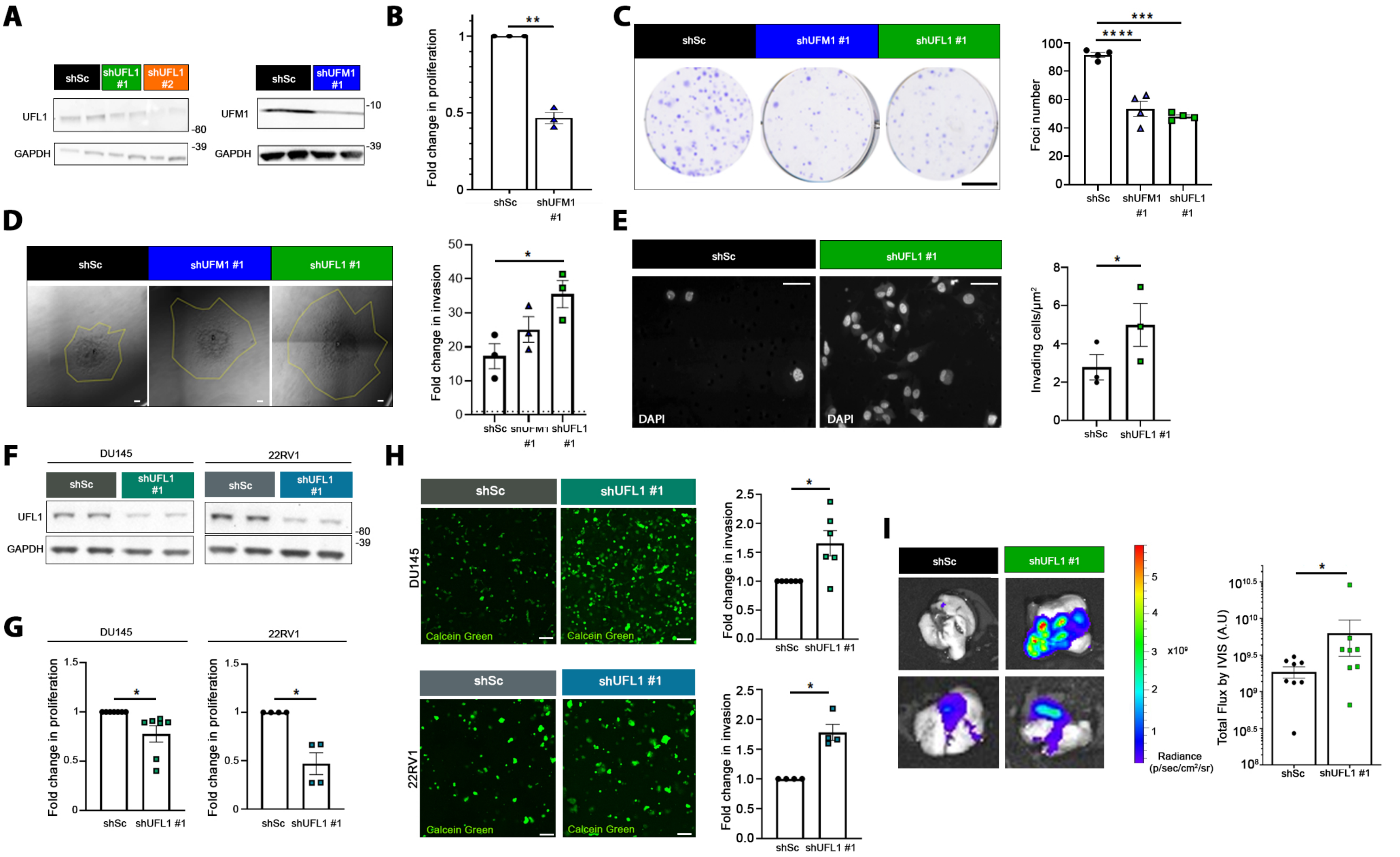

### Extended Data Figure 3

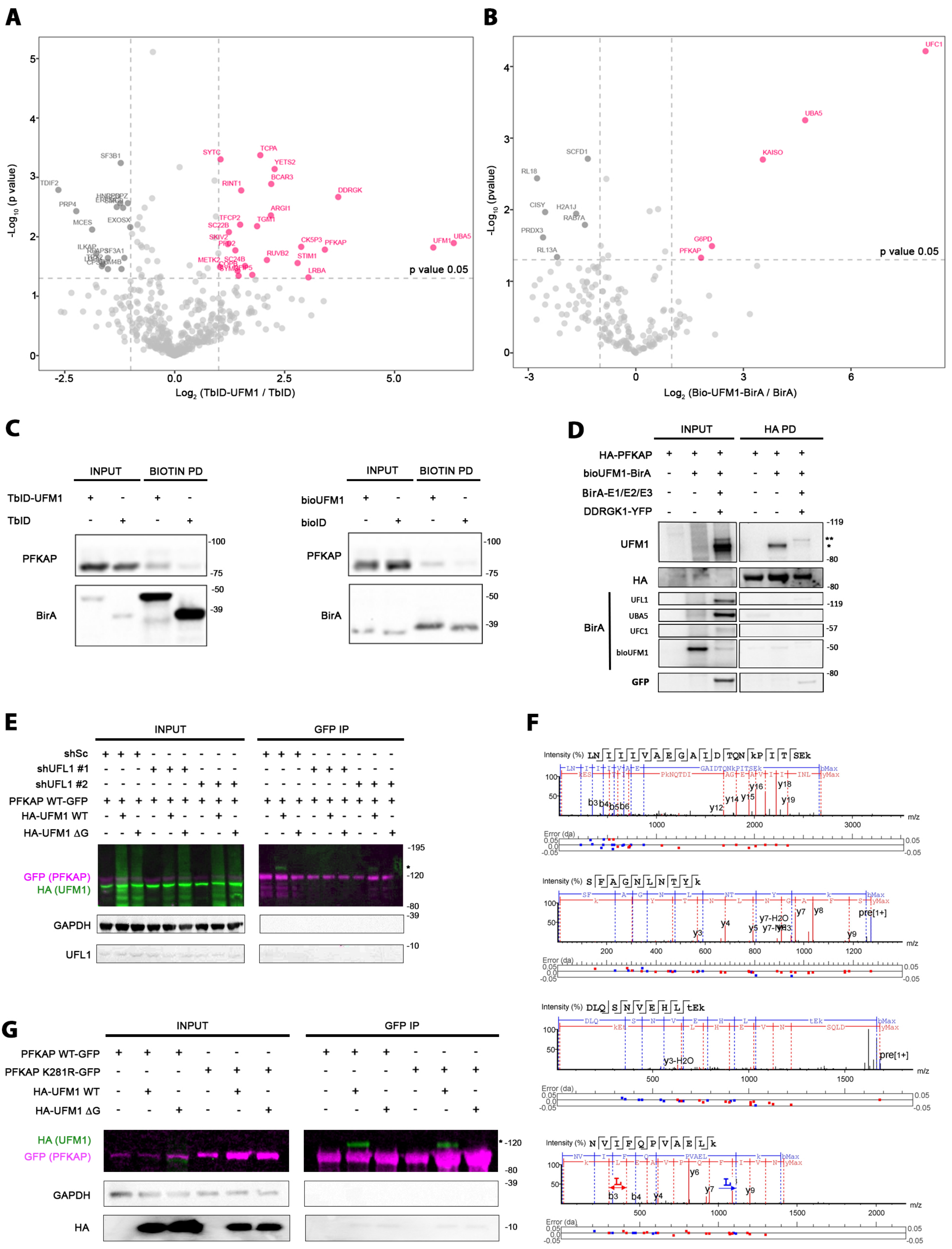

### Extended Data Figure 4

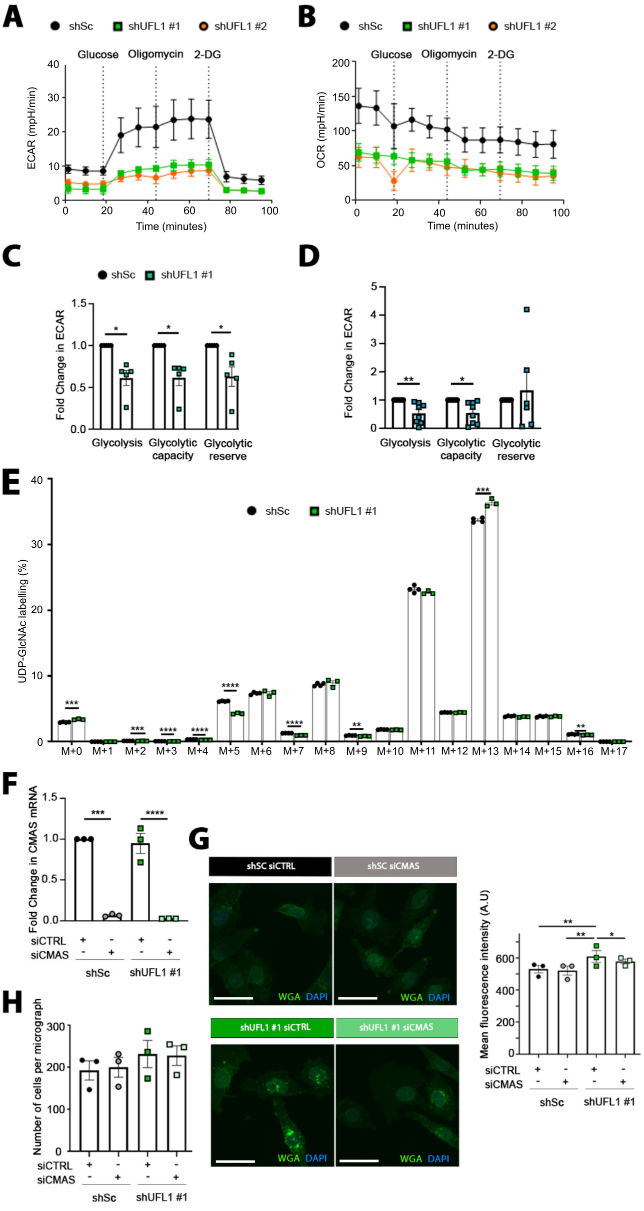

### Extended Data Figure 5

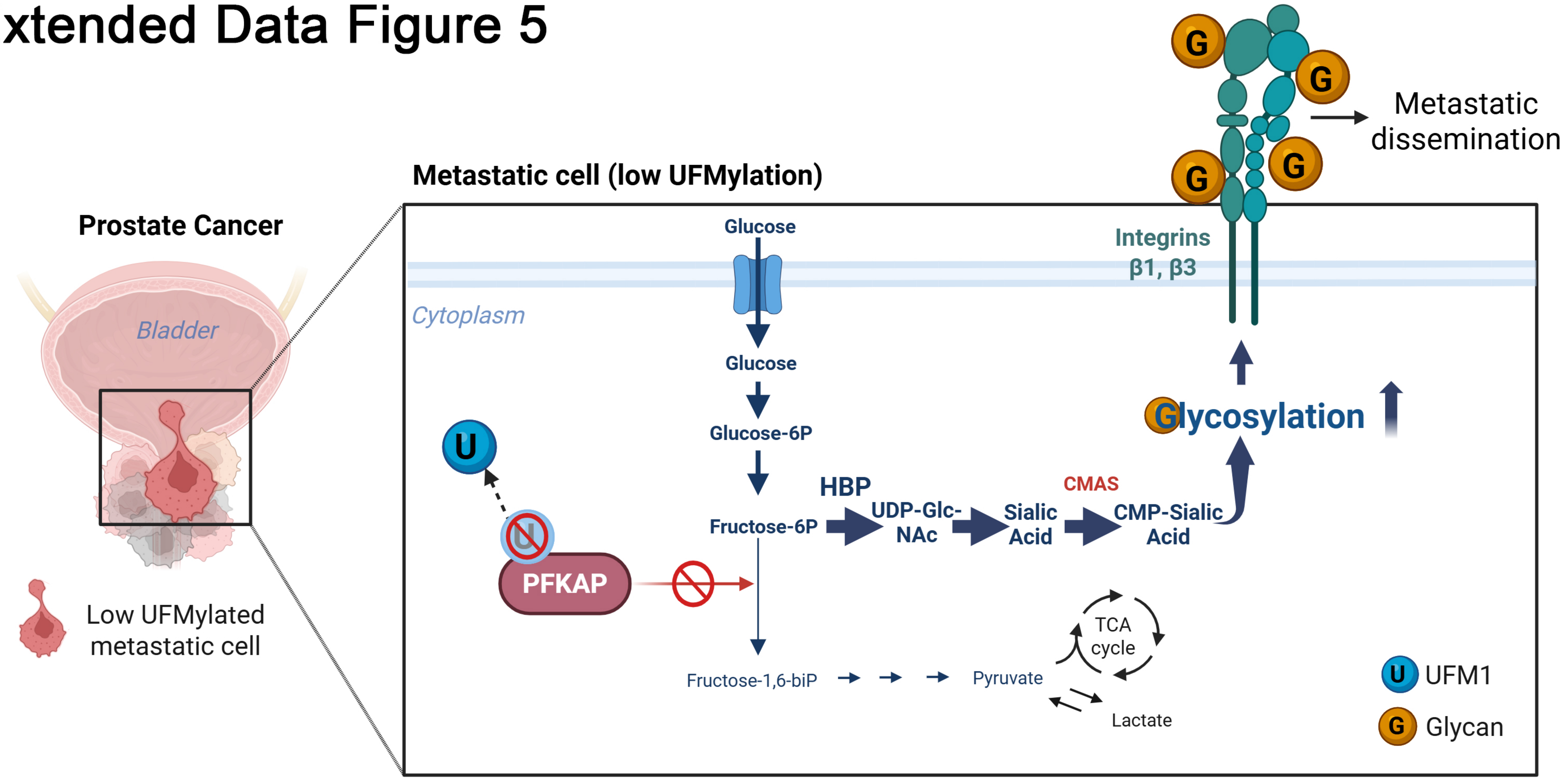

#### EXTENDED DATA FIGURE LEGENDS

**Extended Data Fig. 1. Related to Fig. 1. Copy number aberrations influence *UFL1* and *UFM1* gene expression in prostate cancer.**

**A.** Violin plots depicting the $Log_2$ (mRNA expression) of *UFL1* (upper panels) and *UFM1* (lower panels) in normal (N) and primary tumor (PT) human prostate cancer specimens in the indicated datasets. P value derives from unpaired t-test analysis between the indicated groups. *Source: cancertool.org.*

**B and C.** *UFL1* (B) and *UFM1* (C) $Log_2$ (mRNA expression) sorted by copy number aberrations in prostate cancer specimens. Deep deletion indicates a deep loss, possibly a homozygous deletion; shallow deletion indicates a shallow loss, possibly a heterozygous deletion; gain indicates a low-level gain (a few additional copies, often broad); amplification indicate a high-level amplification (more copies, often focal). *Source: cbioportal.org.*

**D.** Schematic overview of the UFMylation pathway. The precursor form of UFM1 (pro-UFM1) undergoes cleavage by UFSPs to reveal its C-terminal conserved Gly residue (G). Subsequently, UFM1 is activated by UBA5 (activating enzyme E1) through an ATP-consuming process, forming a high-energy thioester bond with UBA5. The E2 conjugating enzyme UFC1 then interacts with UBA5 to release UFM1 and create a thioester bond with UFM1. Finally, UFC1, in conjunction with UFL1 (E3 ligase), transfers UFM1 to its target substrate. DDRGK1 acts as an adaptor protein enabling UFL1 to recruit a broader range of substrates. Furthermore, as UFMylation is reversible, UFSPs can detach UFM1 molecules from their targets. K: lysine residue of the substrate. *Figure created with BioRender.com.*

**Extended Data Fig. 2. Related to Fig. 2. Silencing of *UFM1* and *UFL1* reduces cell growth and promotes invasion and metastatic dissemination of prostate cancer cells *in vitro* and *in vivo.***

**A.** Representative western blot of total lysates of shSc, shUFL1 #1 and shUFL1 #2 PC3 cells. Specific antibodies against UFL1, UFM1 and GAPDH as loading control were used. Molecular weight markers (kDa) are shown to the right.

**B.** Analysis of cell proliferation by crystal violet staining upon *UFM1* silencing (shUFM1 #1). The fold change in crystal violet absorbance at 6 days relative to scramble (shSc) is represented. P value was obtained by one-sample t-test. shSc *vs.* shUFM1 #1 (n=3).

**C.** Representative images (left) and quantification analysis of foci formation in control (shSc), *UFM1*- (shUFM1 #1) and *UFL1*-silenced (shUFL1 #1) PC3 cells. Each dot represents the mean number of foci per condition (n=4). P value was obtained by ordinary one-way ANOVA with multiple comparison analysis. Scale bar: 1 mm.

**D.** Representative images (left) and quantification (right) of the invasive ability of control (shSc), *UFM1*- (shUFM1 #1) and *UFL1*-silenced (shUFL1 #1) PC3 cells in a collagen 3D matrix. The area of the spheroids was measured at day 0 and day 5 and the fold change in invasion at 5 days relative to their respective area at day 0 (dashed line) is represented (n=3). P value was obtained by ordinary one-way ANOVA with multiple comparison analysis. Scale bar: 10 μm.

**E.** Representative images of invading cells in a Matrigel-coated transwell assay are shown (left). Cells were detected by DAPI (white). Scale bar, 10 μm. Graph representing the fold change in number of invading cells per membrane area ($\mu m^2$) normalized to control condition (shSc). P value was obtained by one-tailed paired t-test.

**F.** Representative western blot of total lysates of shSc and shUFL1 #1 DU145 and 22RV1 cells. Specific antibodies against UFL1 and GAPDH as loading control were used. Molecular weight markers (kDa) are shown to the right.

**G.** Analysis of cell proliferation by crystal violet staining of control (shSc) and *UFL1*-silenced (shUFL1 #1 and shUFL1 #2) DU145 (left) and 22RV1 (right) cells. The fold change in crystal violet absorbance at 6 days normalized to shSc is represented. P value was obtained by one-sample t-test. DU145 shSc *vs.* shUFL1 #1 (n=7); 22RV1 shSc *vs.* shUFL1 #1 (n=4).

**H.** Representative images of the invasive ability of control (shSc) and *UFL1*-silenced (shUFL1 #1) DU145 (top) and 22RV1 (bottom) cells in a collagen:matrigel 3D matrix. The invasive area of cancer cells was visualized by calcein green staining. Scale bar: 100 µm. Graphs on the right represent the quantification of the invasive ability of control (shSc) and *UFL1*-silenced (shUFL1 #1) DU145 (top) and 22RV1 (bottom) cells in a collagen:matrigel 3D matrix. The total area occupied by calcein green-stained cells was quantified using ImageJ (DU145: n=6 and 22RV1: n=4). P value was obtained by one-sample t-test.

**I.** *Ex vivo* representative images of lungs from 2 different mice (left panels) and IVIS signal quantification of lung metastasis (right graph) are shown. Each dot represents a different mouse (n=8 mice per group). P value was obtained by one-tailed Mann Whitney test.

**Extended Data Fig. 3. Related to Fig. 3. Validation of PFKAP UFMylation**

**A.** Volcano plot representing the distribution of the candidates identified by proximity proteomics in three independent experiments. Proteins with more than 1-fold change in TbID-UFM1 intensity with respect to the TbID (($Log_2$ (TurboID-UFM1 intensity/TurboID intensity) $\geq$ 0)) and P value < 0.05 ((-$Log_{10}$ (p value) > 1.12)) were considered as UFM1-associated candidates (pink dots).

**B.** Volcano plot representing the distribution of the candidates identified by bioUFM1 technique and proteomics in three independent experiments. Proteins with more than 1-fold change in bioUFM1 intensity with respect to the BirA (($Log_2$ (bioUFM1 intensity/BirA intensity) ≥ 0)) and P value < 0.05 ((-$Log_{10}$ (p value) > 1.12)) were considered as UFMylation targets (pink dots).

**C.** Western blot validation of proteomics data in 22RV1 cells transfected with TbID-UFM1 or TbID alone (left panels) and bioUFM1 or BirA alone (right panels). Specific antibodies (PFKAP and BirA) were used as indicated. Anti-BirA antibody detected the fusion form of TbID-UFM1 (46 kDa) and TbID or BirA (37 kDa). PFKAP antibody detected endogenous PFKAP. Molecular weight markers are shown to the right. PD: pulldown.

**D.** Western blot analysis for validating the UFMylation of exogenous PFKAP. HEK293FT cells were transfected with HA-PFKAP alone or together with functional UFM1 (bioUFM1-BirA) with or without the UFMylation machinery (E1/E2/E3 and DDRK1-YFP). PFKAP was immunoprecipitated with HA agarose beads and blots were probed with the indicated antibodies. * and ** mark the bands of potential mono-UFMylated and poly-UFMylated PFKAP, respectively. Molecular weight markers (kDa) are shown to the right. PD: pulldown.

**E.** Western blot analysis for validating that the UFMylation of exogenous PFKAP is reduced in *UFL1*-silenced cells. HEK293FT cells were transfected with shSc, shUFL1 #1 or shUFL1 #2 together with PFKAP-GFP alone with and without functional UFM1 (HA-UFM1 WT) or defective UFM1 (HA-UFM1 ΔG). PFKAP was immunoprecipitated with GFP beads and blots were probed with the indicated antibodies. UFMylated PFKAP is shown in green (HA) and unmodified PFKAP-GFP in purple (GFP). Molecular weight markers (kDa) are shown to the right. * marks the band of UFMylated PFKAP. IP: immunoprecipitation.

**F.** Identification of lysine residues of PFKAP modified by UFM1 through mass spectrometry: K281 and K287, K395, K625 and K736, in descendent order. GFP was immunoprecipitated from cell lysates of HEK293FT cells expressing PFKAP-GFP and HA-UFM1 WT, followed by mass spectrometry analysis.

**G.** UFMylation assay of the immunoprecipitation of wild-type PFKAP (PFKAP WT-GFP) or its mutant form where one lysine was mutated to arginine (PFKAP K281R-GFP) in HEK293FT cells expressing functional UFM1 (HA-UFM1 WT) or defective UFM1 (HA-UFM1 ΔG). UFMylated PFKAP is shown in green (HA) and unmodified PFKAP-GFP in purple (GFP). Molecular weight markers (kDa) are shown to the right. * marks the band of UFMylated PFKAP. IP: immunoprecipitation.

**Extended Data Fig. 4. Related to Fig. 4. A reduction in UFMylation promotes a metabolic switch towards glycosylation**

**A and B.** Graphical quantification of extracellular acidification rate (ECAR, A) and oxygen consumption rate (OCR, B) along time (min) in a Glycolysis Stress test in control (shSc) and *UFL1*-silenced (shUFL1 #1) PC3 cells by Seahorse analysis (n=5). 2-DG: 2-Deoxy-D-glucose.

**C.** Graphical quantification of extracellular acidification rate (ECAR) -derived parameters of the Glycolysis Stress test in control (shSc) and *UFL1*-silenced (shUFL1 #1) DU145 cells by Seahorse analysis (n=5). P value was obtained by one-sample t-test.

**D.** Graphical quantification of extracellular acidification rate (ECAR) -derived parameters of the Glycolysis Stress test in control (shSc) and *UFL1*-silenced (shUFL1 #1) 22RV1 cells by Seahorse analysis (n=6). P value was obtained by one-sample t-test.

**E.** Graph representing the fractional contribution of $^{13}C_6$-glucose to UDP-GlcNac carbon labelling upon 24h with $^{13}C_6$-glucose, analyzed by LCMS in control (shSc) and *UFL1*-silenced (shUFL1 #1) PC3 cells. M+n represents the carbon number that was labelled by glucose. P value was obtained by multiple unpaired t-test ( shSc: n=4 and shUFL1 #: n=3).

**F.** Graphical representation of *CMAS* mRNA levels detected by qRT-PCR in control (shSc) and *UFL1*-silenced (shUFL1 #1) PC3 cells with functional (siCTRL) or inactive (siCMAS) sialic acid pathway (n=3). P value was obtained by one-sample t-test.

**G.** Representative images (left) and quantification of the levels of β-1,4-GlcNAc and sialic acid-linked residues using wheat germ agglutinin (WGA) staining the cytoplasm in control (shSc) and *UFL1*-silenced (shUFL1 #1) PC3 cells with functional (siCTRL) or inactive (siCMAS) sialic acid pathway. Each dot represents a different experiment (n=3). Green, WGA (β-1,4-GlcNAc- and sialic acid-linked proteins); blue, DAPI nuclear staining. P value was obtained by ordinary one way ANOVA with multiple comparison analysis. Scale bar 30 μm.

**H.** Graphical representation of the number of cells quantified in Figure 4G in control (shSc) and *UFL1*-silenced (shUFL1 #1) PC3 cells with functional (siCTRL) or inactive (siCMAS) sialic acid pathway. P value was obtained by repeated measures ANOVA with multiple comparison analysis. Each dot represents the average number of cells per micrograph in 3 different experiments.

**Extended Data Fig. 5. Schematic of the mechanism by which UFMylation-low cells increase metastatic dissemination from the primary tumour.** GlcNAc, N acetylglucosamine; HBP, hexosamine biosynthesis pathway. The illustration was created using *BioRender.com.*
